# Effect of naturally-occurring mutations on the stability and function of cancer-associated NQO1: comparison of experiments and computation

**DOI:** 10.1101/2022.09.30.510258

**Authors:** Juan Luis Pacheco-Garcia, Matteo Cagiada, Kelly Tienne-Matos, Eduardo Salido, Kresten Lindorff-Larsen, Angel L. Pey

## Abstract

Recent advances in DNA sequencing technologies are revealing a large variability in the human genome. Our capacity to establish genotype-phenotype correlations in such large-scale is, however, limited. This task is particularly challenging due to the multifunctional nature of proteins. Here we describe an extensive analysis of the stability and function of naturally-occurring variants (found in the COSMIC and gnomAD databases) of the cancer-associated human NAD(P)H: quinone oxidoreductase 1 (NQO1). First, we performed *in silico* saturation mutagenesis studies (>5000 substitutions) aimed to identify regions in NQO1 important for stability and function. We then experimentally characterized twenty-two naturally-occurring variants in terms of protein levels during bacterial expression, solubility, thermal stability and coenzyme binding. These studies showed a good overall correlation between experimental analysis and computational predictions; also the magnitude of the effects of the substitutions are similarly distributed in variants from the COSMIC and gnomAD databases. Outliers in these experimental-computational genotype-phenotype correlations remain, and we discuss these on the grounds and limitations of our approaches. Our work represents a further step to characterize the mutational landscape of NQO1 in the human genome and may help to improve high-throughput *in silico* tools for genotype-phenotype correlations in multifunctional proteins associated with disease.

## 1. Introduction

Advances in technologies for whole-genome or exome sequencing and high-throughput functional assays have increased our knowledge on the consequences of the genetic variability in humans and the relationship to disease (1–4). However, our capacity to predict the pathogenicity of single amino acid variants is still limited, with some approaches providing good overall results but failing to predict correlation for some individual mutations or phenotypes (3).

Current approaches for correlating genotype and phenotype can broadly be classified into two classes. First, experimental approaches based on the characterization of one or several functional features (for example enzymatic function and regulation, protein-protein interactions, transport to different intracellular or extracellular locations, protein turnover, ligand binding) (2,5–7). In the case of high-throughput experimental approaches typically only one or two aspects of protein function are analysed (for example protein abundance or ability to rescue a growth phenotype) (8). Second, the use of structure- or sequence-based methods to predict pathogenicity are becoming increasingly robust (1,2,7). Although experiments may be implemented in a high-throughput fashion, it has until now been limited to a relatively small set of proteins and assays (1,2). Thus, while computational approaches also have limitations, they may be appealing due to their potential application on a proteomic scale (1,2).

In this work, we apply both types of approaches to increase our understanding of the correlation between genotype and phenotype for missense variants of the human NAD(P)H:quinone oxidoreductase 1 (NQO1) protein. NQO1 is associated with several diseases including cancer, Alzheimer’s and Parkinson’s disease (9,10). NQO1 is a multifunctional protein, displaying both enzymatic and non-enzymatic functions. As an enzyme, it catalyzes the FAD-dependent (two-electron) reduction of a large set of quinone substrates, with functions including redox maintenance of vitamins, detoxification of xenobiotics and activation of cancer pro-drugs (10–13). Among non-enzymatic functions, NQO1 may interact with proteins and RNA, controlling their stability and function (10,14–17). Many of these functions are associated with the catalytic competence and FAD binding, such as protein-protein interactions, intracellular stability and association with microtubules (14,18–20). The native form of NQO1 is dimeric, containing two different domains: an N-terminal domain (NTD, residues 1-225), that tightly binds one FAD molecule/domain required for catalysis and contains most of the monomer-monomer interface (MMI), whereas the C-terminal domain (CTD, residues 225-274) complete the active site and the MMI (5,21–24).

We have recently shown that ligand binding and variant effects on stability propagate to long distances in the native state, affecting different functional features in a counterintuitive fashion (5,25–30). Therefore, NQO1 represents a challenging and biomedically relevant system to compare the performance of computational and experimental methods to explain and to predict genotype-phenotype in a large-scale for a multi-functional protein. Here, we use computational tools to probe 5187 variants of NQO1 that includes a set of clinically relevant missense variants which we then experimentally characterized. In this set, thirteen variants come from large-scale human sequencing data (gnomAD) and nine from the catalogue of somatic mutations in human cancer lines (COSMIC) (Table 1). As of 9^th^ of January 2021, none of these variants were found in both databases. Whether variants found in COSMIC or gnomAD databases are associated with disease (e.g. predisposition to cancer development) is unknown. The set of variants we studied experimentally comprises very different amino acid chemistries and display different levels of solvent exposure (Table 1).

**Table 1.** Set of NQO1 variants experimentally characterized in this work. The table indicates whether the variants are found in the COSMIC/gnomAD databases as well as the solvent exposure (as % ASA) determined using a crystallographic model of WT NQO1 (PDB code 2F1O; (54)), the software GetArea and the computational classification at variant and residue level using a combination of predictions of thermodynamic stability change upon mutation and evolutionary conservation.

| **Mutation** | **Database** | **% ASA\*** | **Variant Class** | **Residue Class** |
| --- | --- | --- | --- | --- |
| G3S | gnomAD | 6.0±4.7 | WT-like | WT-like |
| G3D | COSMIC | 6.0±4.7 | WT-like | WT-like |
| L7P | COSMIC | 0.3±0.2 | Total-loss | Total-loss |
| L7R | gnomAD | 0.3±0.2 | Total-loss | Total-loss |
| V9I | gnomAD | 0.0±0.0 | WT-like | Total-loss |
| T16M | gnomAD | 43±14 | Stable but inactive | WT-like |
| Y20N | gnomAD | 21±5 | WT-like | WT-like |
| A29T | COSMIC | 2.2±0.5 | WT-like | Unstable but active |
| K32N | gnomAD | 79±11 | WT-like | WT-like |
| G34V | gnomAD | 54±11 | Total-loss | Stable but Inactive |
| E36K | gnomAD | 64±3 | WT-like | WT-like |
| S40L | gnomAD | 0.0±0.0 | WT-like | Total-loss |
| D41G | gnomAD | 14±2 | Total-loss | Stable but inactive |
| D41Y | COSMIC | 14±2 | Stable but inactive | Stable but inactive |
| M45L | COSMIC | 28±5 | WT-like | WT-like |
| M45I | COSMIC | 28±5 | WT-like | WT-like |
| I51V | gnomAD | 14±1 | WT-like | Stable but inactive |
| W106R | gnomAD | 10±1 | Total-loss | Total-loss |
| W106C | COSMIC | 10±1 | Total-loss | Total-loss |
| F107C | gnomAD | 7.6±0.5 | Unstable but | Unstable but |
|  |  |  | active | active |
| M155I | COSMIC | 13±3 | WT-like | WT-like |
| H162N | COSMIC | 6.6±0.5 | WT-like | WT-like |

\* Using GetArea (http://curie.utmb.edu/getarea.html) and the structure with PDB code 2F1O (54). Data are the average ± s.d. from eight monomers.

## 2. Materials and Methods

### 2.1 Experimental methods

#### 2.1.1 Protein expression and purification

Mutations were introduced by site-directed mutagenesis in the wild-type (WT) NQO1 cDNA cloned into the pET-15b vector (pET-15b-NQO1) by GenScript (Leiden, Netherlands). Mutated codons were optimized for expression in *E. coli* and mutagenesis was confirmed by sequencing the entire cDNA. The plasmids were transformed in *E. coli* BL21(DE3) cells (Agilent Technologies, Santa Clara, CA, USA) for protein expression.

To determine the amount of soluble NQO1 at 37 °C, 5 mL of LB medium containing 0.1 mg·mL^−1^ ampicillin (purchased from Canvax Biotech, Córdoba, Spain) was inoculated with transformed cells and grown for 16 h at 37 °C. 0.5 mL of these cultures were diluted into 10 mL of LB containing 0.1 mg·mL^−1^ ampicillin (LBA) and grown at 37 °C for 3 h. After that, cultures were induced with 0.5 mM of isopropyl β-D-1-thiogalactopyranoside (IPTG, Canvax Biotech) at 37 °C for 4 h. Cells were harvested by centrifugation at 2900 *g* in a bench centrifuge at 4 °C and frozen at −80 °C for 16 h. Cells were resuspended in binding buffer (20 mM Na-phosphate, 300 mM NaCl, 50 mM imidazole, pH 7.4) with 1 mM phenylmethylsulfonyl fluoride (PMSF, Sigma-Aldrich, Madrid, Spain) and sonicated in an ice bath. These *total extracts* were centrifugated (24000 *g*, 30 min, 4 °C in a bench centrifuge) to obtain the *soluble extracts*. The amount of NQO1 in total and soluble extracts was determined by Western-blotting providing the S/T (soluble/total) ratio for each variant. Samples were denatured using Laemmli’s buffer, resolved in polyacrylamide gel electrophoresis in the presence of sodium dodecylsulphate (SDS-PAGE, 12% acrylamide) gels and transferred to PVDF membranes (GE Healthcare, Chicago, IL, USA) using standard procedures. Immunoblotting was carried out using primary monoclonal antibody against NQO1 (sc-393736, Santa Cruz Biotechnology, Dallas, TX, USA) at 1:500 dilution and, as secondary antibody, an anti-mouse IgGκ BP-HRP (sc-516102, Santa Cruz Biotechnology) at 1:2000 dilution was used. Samples were visualized using luminol-based enhanced chemiluminescence (from BioRad Laboratories, Hercules, CA, USA), scanned and analysed using Image Lab (from BioRad Laboratories).

For large-scale purifications, a preculture (100 mL) was prepared from a single clone for each variant and grown for 16 h at 37 °C in LBA and diluted into 2.4-4.8 L of LBA. After 3 h at 37 °C, NQO1 expression was induced by the addition of 0.5 mM IPTG for 6 h at 25 °C. Cells were harvested by centrifugation at 8000 *g* and frozen overnight at −80 °C. NQO1 proteins were purified using immobilized nickel affinity chromatography columns (IMAC, GE Healthcare) as described (12). Isolated dimeric fractions of NQO1 variants were exchanged to HEPES-KOH buffer 50 mM pH 7.4 using PD-10 columns (GE Healthcare). The UV–visible spectra of purified NQO1 proteins were measured in a Cary spectrophotometer (Agilent Technologies, Waldbronn, Germany) and used to quantify NQO1 concentration and the content of FAD as described in (12). Apo-proteins were obtained by treatment with 2 M urea and 2 M KBr as described in (27), obtaining samples with less than 2% bound FAD based on UV-visible spectra. Samples were stored at −80 °C upon flash freezing in liquid N_2_. Protein purity and integrity was checked by SDS-PAGE.

#### 2.1.2 In vitro characterization of NQO1 variants

Thermal denaturation of NQO1 proteins, as holo-proteins (2 μM in monomer +100 μM FAD) was monitored by following changes in tryptophan emission fluorescence in HEPES-KOH 50 mM at pH 7.4 as described in (31). *T*_m_ values were reported as mean ± s.d. of four independent measurements.

Fluorescence titrations were carried out at 25 °C using 1 × 0.3 cm path-length cuvettes in a Cary Eclipse spectrofluorimeter (Agilent Technologies, Waldbronn, Germany). Experiments were carried out in 20 mM K-phosphate, pH 7.4, essentially as described in (25). Briefly, apo-NQO1 (0.25 μM in subunit) was mixed with 0–2 μM FAD in K-phosphate 20 mM pH 7.4. Samples were incubated at 25 °C in the dark for at least 10 min before measurements. Spectra were acquired in the 340–360 nm range upon excitation at 280 nm (slits 5 nm), and spectra were averaged over 10 scans registered at a scan rate of 200 nm·min^−1^. FAD binding fluorescence intensities at 350 nm were fitted using a single and identical type of binding sites as described in (25). Variant effects on the FAD binding free energy (ΔΔG_FAD_) were calculated as:

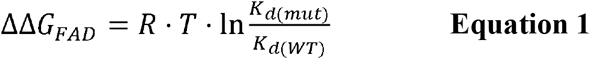

Where R is the ideal gas constant (1.987 cal·mol·K^−1^), T is the experimental temperature (298.15 K), and *K*_d(mut)_ and *K*_d(WT)_ are the FAD binding dissociation constant of the mutant and WT protein variants, respectively. A positive value of ΔΔG_FAD_ indicates that the mutation reduces the affinity for FAD.

For proteolysis studies, NQO1 samples (10 μM in monomer) were prepared in HEPES-KOH 50 mM at pH 7.4 in the presence of 100 μM FAD (NQO1_holo_) and incubated at 25°C for 5 min. The proteolysis reaction was initiated upon addition of 0.02–1.2 μM thermolysin (from *Geobacillus stearothermophilus*, Sigma-Aldrich) and a final concentration of 10 mM CaCl_2_. Samples were incubated at 25 °C and aliquots were collected over time and the reaction quenched by addition of EDTA pH 8 (final concentration of 20 mM) and Laemmli’s buffer (2x). Controls (time 0) were prepared likewise but without thermolysin. Samples were resolved by SDS-PAGE under reducing conditions in gels containing 12% acrylamide. Gels were stained with Coomassie blue G-250. Densitometry was carried out using ImageJ. Data were analyzed using an exponential function to provide the apparent rate constant (*k*_obs_). From the linear dependence of *k_obs_* vs. [thermolysin], we obtained the second-order rate constant *k*_prot_. Linearity in these plot indicate that proteolysis occurs under a EX2 mechanism, thus reflecting the thermodynamic stability of the thermolysin cleavage site (Ser72-Val73) between non-cleavable and cleavable states (32). These *k*_prot_ values were used to determine mutational effects on the local stability of thermolysin cleave site (ΔΔG_PROT_) using equation 2:

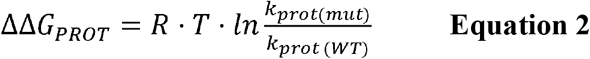

Where R is the ideal gas constant (1.987 cal·mol·K^−1^), T is the experimental temperature (298.15 K), and *k*_prot(mut)_ and *k*_prot(WT)_ are the second-order proteolysis rate constants of the mutant and WT protein variants, respectively. A positive value of ΔΔG_PROT_ indicates that the mutation thermodynamically destabilizes the thermolysin cleavage site.

### 2.2. Computational analyses

#### 2.2.1 Evolutionary Conservation analysis

We used GEMME (33) to evaluate evolutionary distances from the WT NQO1 sequence (Uniprot ID: P15559 — isoform 1). We used HHblits (34,35) to generate a multiple sequence alignment (MSA) using UniClust30 as sequence database and an E-value threshold of 10^−10^. The resulting MSA contained 1602 sequences and was refined using two additional filters: first, all the columns that were not present in the WT NQO1 sequence were removed; second, all the sequences with more than the 50% of gaps were removed. Application of these two filters yielded 1414 sequences in the MSA. The GEMME package was run using default parameters. For each position, a median score was evaluated using all the available substitutions.

#### 2.2.2. Thermodynamic stability predictions

Changes in thermodynamic stability (ΔΔG) were calculated using the crystal structure (21) (PDB 1D4A) and the Rosetta package (GitHub SHA1 c7009b3115c22daa9efe2805d9d1ebba08426a54) with the Cartesian ΔΔG protocol (36,37). The values obtained from Rosetta in internal Energy Unit were divided by 2.9 to bring them on to a scale corresponding to kcal·mol^−1^ (37). Median scores were evaluated for each position using all the available substitutions.

We used DSSP (38) to calculate the solvent accessible surface area (SASA) when identifying interface residues in NQO1. Interface residues were defined as those residues for a difference larger than 0.2 was detected between SASA calculations based on the dimer and monomer structure.

## 3. Results

### 3.1 Saturation mutagenesis by computational methods

We first used the predictive ability of evolutionary conservation analysis combined with thermodynamic stability calculations to classify all possible variants (i.e. saturation mutagenesis) in NQO1 based on their effects on the protein function and stability (8). For evolutionary conservation studies, we used GEMME (33) which provides a score (ΔE) for all possible single amino acid change variants of NQO1 (Figure S1A). ΔE represents the evolutionary distance of a variant from the WT NQO1 sequence, and ΔE has been shown to be a useful predictor of the deleterious effects on function and stability of the given substitution. We used Rosetta (37) to predict variant effects on thermodynamic stability (ΔΔG) using the crystal structure of NQO1 (21) as input (Figure S1B) and subsequently calculated the median ΔΔG for all variants at each position. We performed ΔΔG evaluations using both the monomeric and dimeric structure of NQO1 to separate effects on overall stability and effects on dimerization. Specifically, we calculated ΔΔG from the monomer (Figure S1B and S2A) to predict the change in thermodynamic stability relative to wild type of each variant. We also performed similar ΔΔG calculations using the dimer structure as input (Figure S1C and S2B), introducing each missense variant in both chains (i.e. treating this as a homodimer) and used the resulting values to compare with experiments. Based on these two calculations, we also evaluated the ΔΔG of dimerization as the difference between the two Rosetta runs (Figure S2C and D) to highlight which residues are involved in stabilizing the dimer and thus also those variants that might weaken dimer formation. Then, we evaluated the difference in the SASA between the dimeric and monomeric residues of NQO1 (Figure S2C) and we classified 33 of them as interface residues. We found that for 20 of these interface residues the median ΔΔG of dimerization was larger than 1 kcal·mol^−1^. Of these 20, 15 were stable upon mutation in the monomeric form (median ΔΔG < 2 kcal·mol^−1^) and a subset of 7 display a median ΔΔG of dimerization larger than 2 kcal·mol^−1^.

Then, we combined the evolutionary conservation scores and stability calculations based on the monomeric protein for the 5187 variants of NQO1 and plotted the results in a two-dimensional histogram (Figure 1A). We used cutoff values of 2 kcal·mol^−1^ for Rosetta ΔΔG values and −3 for GEMME ΔE scores as thresholds for all the variants in order to separate them based on their effects (9). To ease analyses and interpretations, we associated each of the four defined regions with a color (8). ‘WT-like’ variants represent 48% of the available NQO1 variants (shown in green). 20% of NQO1 substitutions show high ΔΔG and high evolutionary distances and are classified as ‘Total-loss’. These variants have substitutions that are unlikely in the evolutionary analysis (ΔE < −3) and lead to decreased stability (ΔΔG > 2 kcal·mol^−1^); they thus likely compromise protein function via loss of protein stability (colored in red). Variants with high negative ΔE and low ΔΔG belong to the ‘Stable but inactive’ class (colored in blue). This class contains 24% of the variants and represent those for which the evolutionary and stability analysis suggests that the protein function has been compromised, but not for stability reasons. Lastly, the remaining 8% of the variants show low stability and low evolutionary distance, and were classified as ‘Unstable but active’ (colored in yellow). Having predicted the effects of all possible missense variants, we performed a similar classification of amino acid positions, assigning the most common variant class to each position (Figure 1C-D) and found 48% of the total positions classified as ‘WT-like’, 25% as ‘Total-loss’, 22% as ‘Stable but inactive’ and 5% as ‘Unstable but active’.

**Figure 1.**
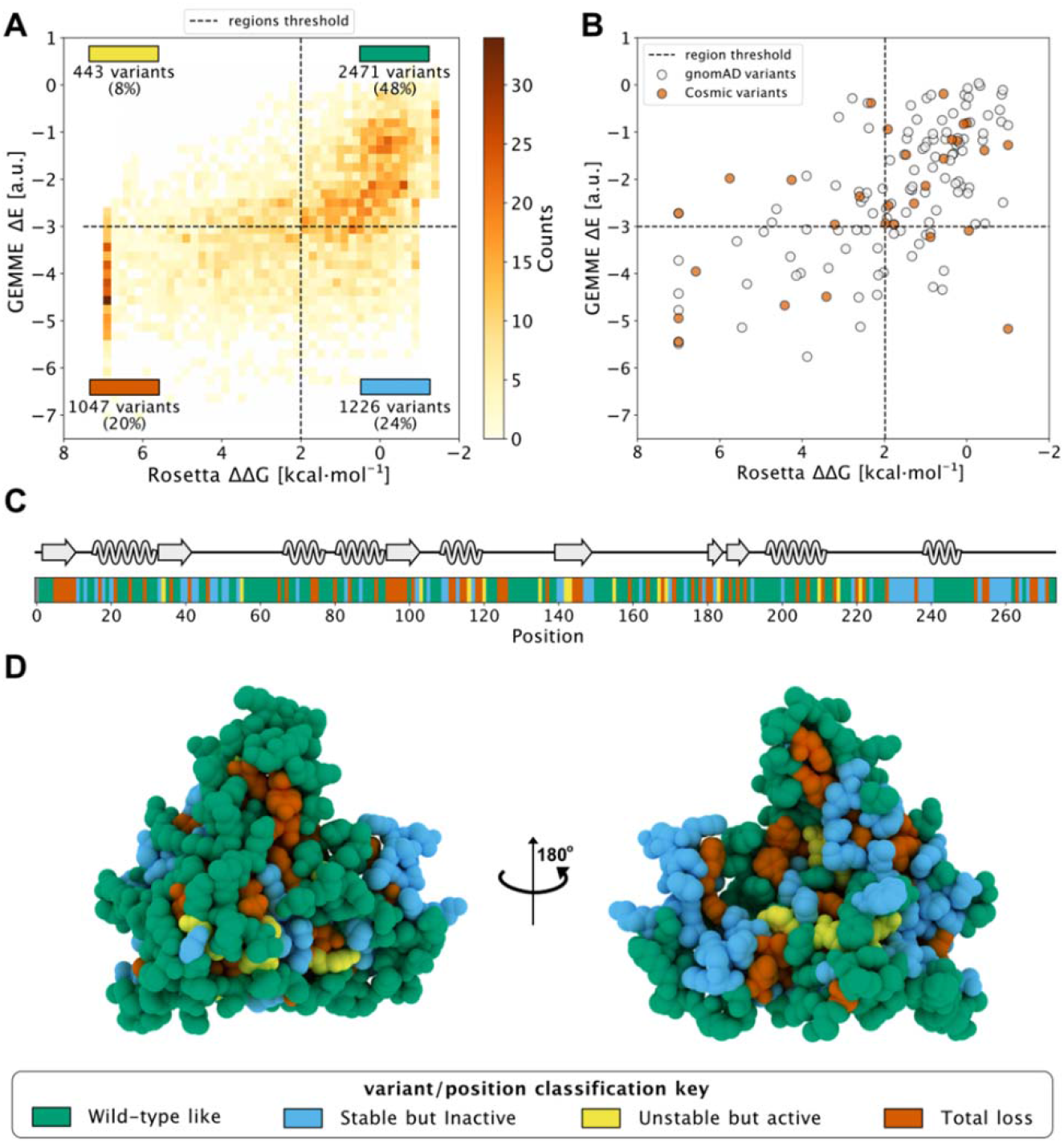
Saturation mutagenesis of NQO1 based on computational methods. A: Two-dimensional histogram which combines the data from the evolutionary conservation analysis (ΔE, y-axis) with the thermodynamic stability (ΔΔG, x-axis) data from Rosetta on the NQQ1 monomer. The variants are categorized in one of the four regions, which are delimited by dashed lines. The fraction of variants, class and colour assigned to each region are indicated. The four classes of variants/positions are reported at the bottom of the figure: “WT-like” (Green), “Low stability, active” (yellow), “Stable but inactive” (blue) and “Total-loss” (red). Arrows on the axes point to region with higher stability or less close to WT in evolutionary space. Panel B reports the scores of gnomAD (grey) and COSMIC (orange) variants in the 2D histogram. Panel C shows the positional colour categories assigned using the most common colour of the position in the sequence together with the secondary structure. Panel D reports the positional classification mapped to the protein crystal structure (PDB: 1D4A)(21).

In addition, we used the data from the dimer analysis to evaluate the number of residues involved in the stabilization of the dimer form. We found that 14 residues at the interface changed their classification to ‘Total-loss’ if ΔΔG was evaluated using the dimer structure. Of these 14, 9 were classified as ‘Stable but inactive’, while 5 were classified as WT-like using monomeric ΔΔG data.

Having analyzed all possible missense variants, we next looked at the results for a subset of variants that have been found in the human population. Specifically, we looked at variants that are found in the COSMIC (COSMIC v.92; https://cancer.sanger.ac.uk/cosmic) or gnomAD (gnomAD v.2.1.1; https://gnomad.broadinstitute.org/) databases, and did not find clear differences between these two sets (Figure 1B). In particular, we found variants in both sets that would be predicted as functional and others for which stability and/or conservation analyses predict loss-of-function (LoF). This result is in line with the notion that both databases may contain both potentially pathogenic as well as benign variants.

### 3.2 Selection of NQO1 variants to be experimentally characterized

After studying the NQO1 variants computationally, we next examined a set of the variants using a series of different experiments. In this study, we have thus extended our previous work on eight naturally-occurring variants in NQO1 (25) to a set of twenty-two variants (Table 1 and Figure 2). Overall, this set included thirteen variants found in the gnomAD database and nine variants found in the COSMIC database. Seventeen of these variants clustered in the N-terminal part of the protein (residues 1–51), whereas five were located in the segment comprising residues 106–162 (in close proximity to the active site). Nine variants affected residues buried inside the protein structure (with less than 10% of SASA), whereas the rest are at positions that are partially or highly solvent-exposed (Table 1). The chemical nature of the changes introduced by the substitutions is also quite diverse, and the substitutions are located in different elements of secondary structure (Figure 2B). Based on our computational analysis the 22 variants represent well the heterogeneous scale of effects on NQO1 function and stability. Indeed, of the 22 variants selected 14 are classified by the computational models as ‘WT-like’, 4 as ‘Total-loss’ and 4 as ‘stable but inactive’.

**Figure 2.**
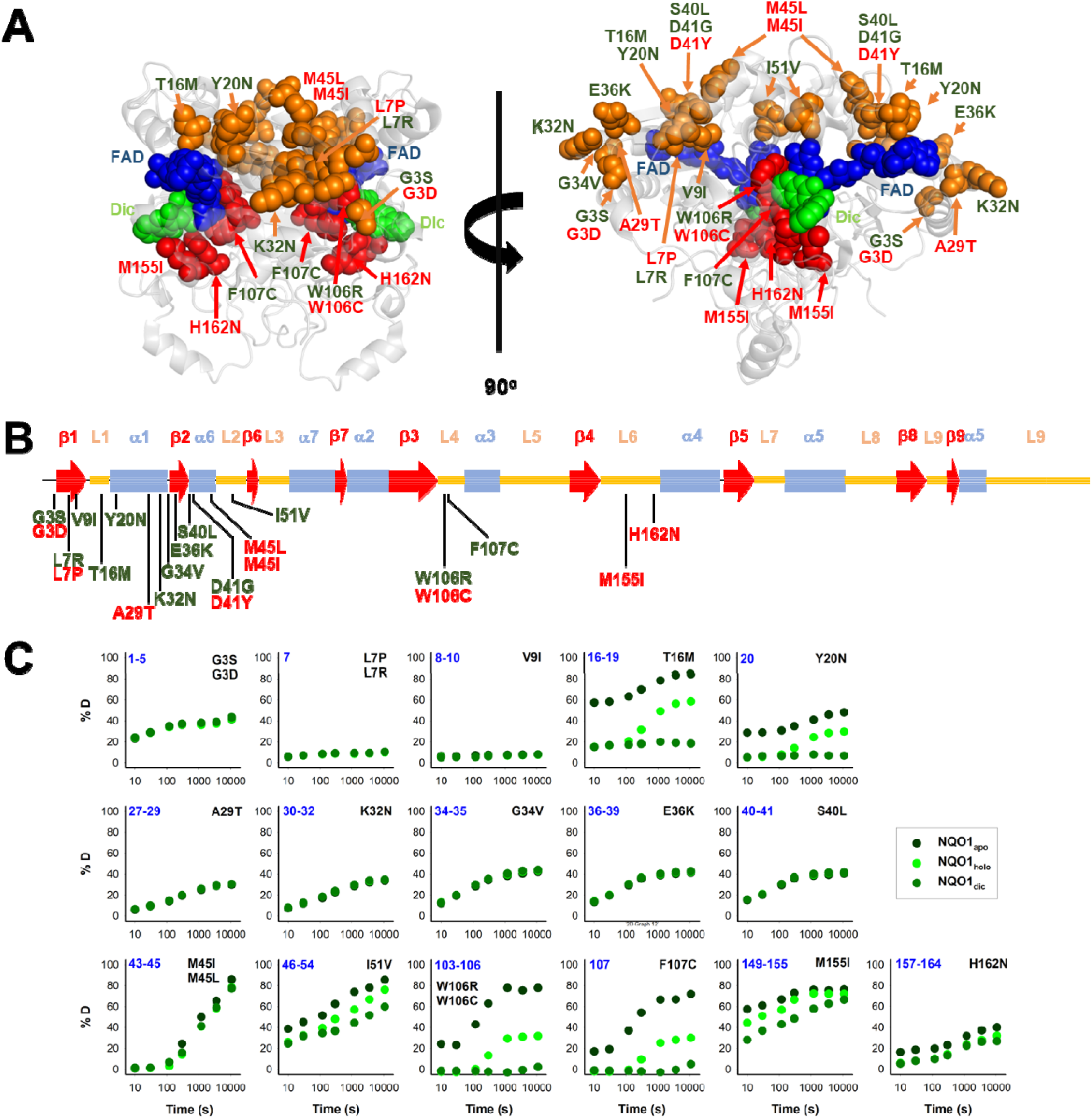
Structural features and local stability of the substituted residues and the variants characterized experimentally in this work. A) The residues mutated were mapped onto the structure of NQO1 (PDB code 2F1O) (54). Residues are depicted as spheres, and the color code indicates substitutions located in the 1-51 region (orange) or the active site (red). Variants were labelled in red (from COSMIC) or in green (from gnomAD). FAD and Dic are shown as dot representations in cyan and blue, respectively. B) Location of substitutions in the sequence of NQO1 regarding secondary structure elements (from (21)). Substitutions were labelled in red (from COSMIC) or in green (from gnomAD). C) HDX of segments containing the 22 variants experimentally investigated in this work. The segments are labelled in blue. The colour code corresponds to HDX for NQO1_apo_, NQO1_holo_ and NQO1_dic_ states. HDX data are from (27).

Results from a recent hydrogen/deuterium exchange (HDX) study on WT NQO1 (27) enables us to evaluate the local stability of the protein segments in which these residues are found as well as the effect of FAD and dicoumarol binding (two ligands of functional and stability relevance) (Figure 2C). The L7P, L7R and V9I substitutions are located in regions with high stability that do not change upon binding of FAD or dicoumarol (Dic; a competitive inhibitor of NADH). Variants G3S, G3D, A29T, K32N, G34V, E36K, S40L and H162N are located in regions with intermediate HDX stability and whose local stability is hardly sensitive to ligand binding. Nevertheless, these positions could still, in principle, affect protein folding, stability, or solubility and indirectly affect the binding of the substrates. M45L and M45I are found in a region with low stability and not responsive to ligand binding. Y20N is located in a region with intermediate stability and where HDX shows a response to ligand binding; the remaining six variants (T16M, I51V, W106R, W106C, F107C and M155I) are in regions with low stability and also their local stability respond directly to ligand binding. This last group of variants may thus have a greater potential to affect enzymatic activity (preventing either the formation of the “stable” holo-protein, a precatalytic state, or the formation of a catalytically relevant state, the Dic state, with FAD and the inhibitor Dic bound; (12)). Although this suggestion is simple, it must be noted that regions of either high or low local stability may play roles in enzyme functional and allosteric response, and that local effects can propagate far from the perturbed site (due to ligand binding or amino acid substitutions) (39–41). Therefore, we performed an experimental characterization of variant effects on protein stability and function and compared the results with our computational analyses.

### 3.3. Variant effects on the expression levels, solubility and stability of NQO1

We first experimentally analysed the effect of these 22 NQO1 variants on the expression levels and solubility of the protein (upon expression in *E.coli*) as well as their effects on conformational stability by thermal denaturation experiments (Table S1 and Figures 3–4).

**Figure 3.**
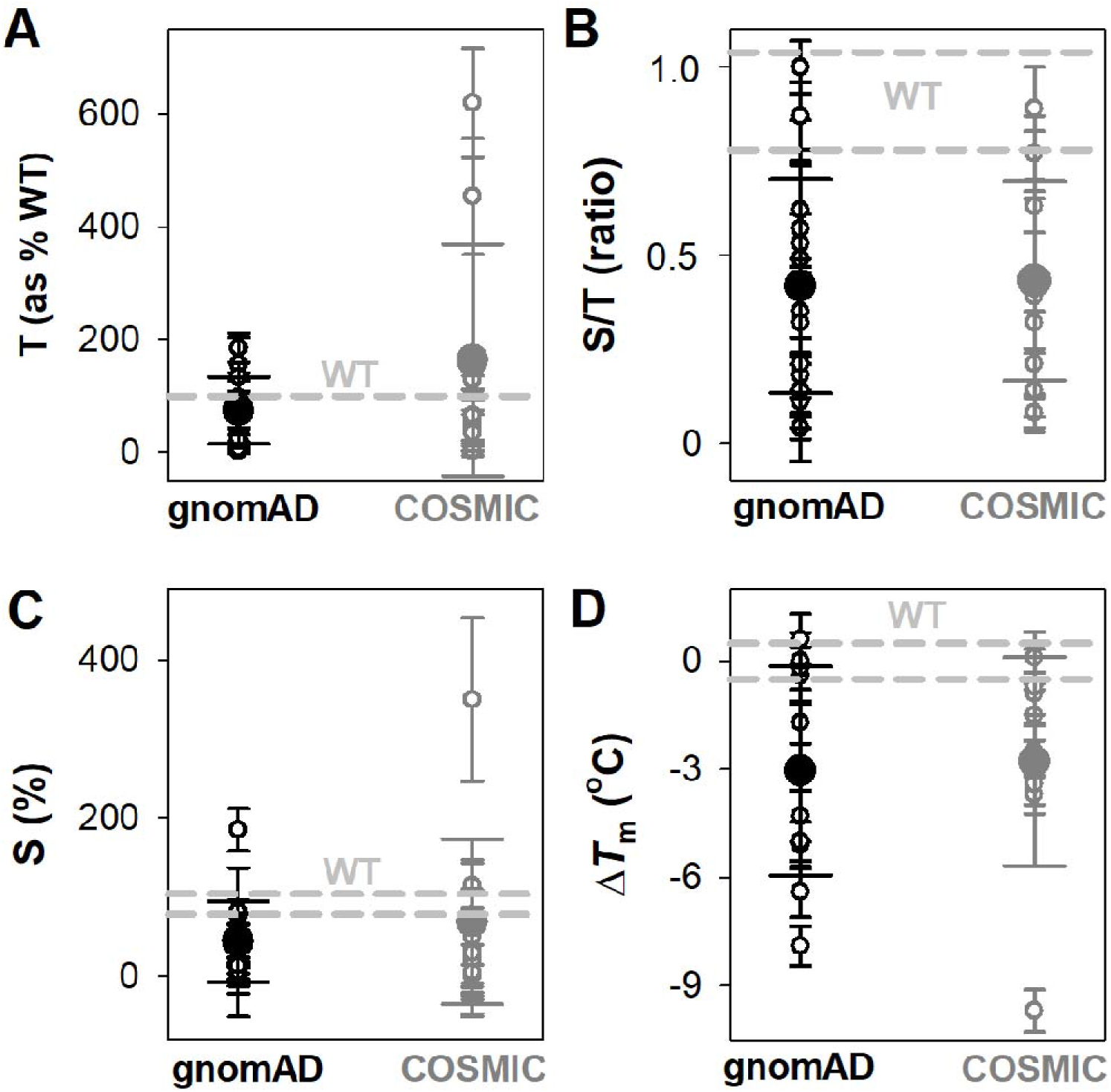
Overall effects of gnomAD and COSMIC variations on the aggregation/solubility and thermal stability of the NQO1 protein. A-C) Expression analyses of aggregation/solubility of NQO1 variants in *E.coli* at 37°C. The total amount of NQO1 protein (T, panel A) as well as the ratio of soluble/total (S/T) protein (B, panel B) were determined from induced cells sonicated before (T) and after (S) centrifugation at 21000 g. The levels of NQO1 protein were determined by western-blotting (Figure S3). The amount of soluble protein (S) is calculated as the product of T and S/T and shown in panel C. Data are the mean ± s.d. from at least three independent expression experiments for each variant.; D) Thermal stability of NQO1 variants as holo-proteins. Δ*T*_m_ values correspond to the difference between a given variant and the WT protein. Data are the mean ± s.d. from three-six technical replicas. Small circles indicate the effects of individual variants and large circles (and corresponding errors) are those for each data set (mean±s.d.). For reference, the values corresponding to WT NQO1 are shown in light grey. Variants are grouped in the gnomAD and COSMIC sets.

**Figure 4.**
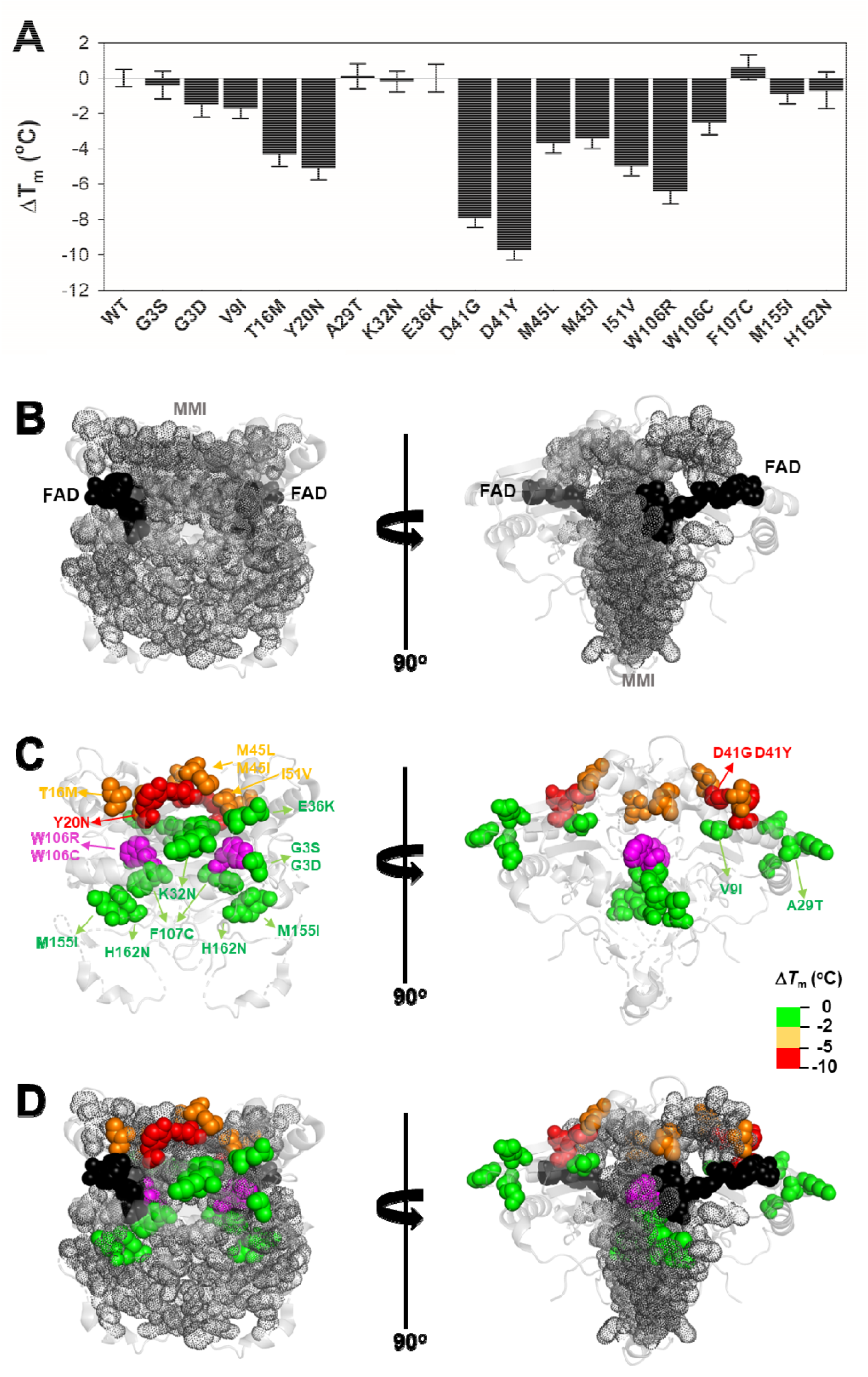
Variant effects on thermal stability related to their location near the MMI. A) Experimental Δ*T*_m_ values for individual variants; B-D) Structural location of the FAD (black spheres) and the MMI (grey dots) (panel B) and mutated residues (color scale according to their destabilizing effect) (panel C). Panel D shows an overlay of panels B and C. Note that two views rotated 90° are shown. The structural model used for display was PDB code 2F1O (54). The residue W106 is highlighted in magenta due to the widely different effects of the W106R/W106C substitutions.

The analysis of the total (T) expression of the variants vs. WT NQO1 at 37°C (Figure 3A-B and S3; Table S1) revealed that some variants (G3S, G3D, L7P and V9I) showed higher total expression levels, in agreement with our previous report (25). This is likely the result of codon optimization used in the mutagenesis. Most of the variants showed relatively high expression levels, ranging from 25% to full WT levels, indicating that these variants mildly to moderately reduced total expression levels. The L7R, G34V, S40L, D41G and D41Y variants showed extremely low expression levels, thus preventing further biophysical characterization. Interestingly, overall, no substantial differences were observed between the average effect of the gnomAD and COSMIC sets of variants.

We then determined the fraction of NQO1 existing as soluble protein (S/T ratios; Figure 3B and Table S1). WT NQO1 showed a ratio of ~ 0.9 (Table S1 and Figure 3B). Again, although some variants showed much lower S/T ratios than WT NQO1, most of them showed values between 0.2 and the WT ratio. Only five variants showed lower S/T ratios than 0.15 (L7P, L7R, S40L, F107C and M155I). Expression under milder conditions (25 °C) did not allow for purification of enough protein of the L7P, L7R, S40L and G34V variants for further biophysical characterization.

We used the product of total expression levels and S/T ratios (i.e. the S values) as a proxy to evaluate the overall effect of amino acid substitutions on NQO1 solubility-aggregation propensity when expressed at 37 °C (Table S1 and Figure 3C). Nine substitutions reduced the solubility below 20% of the WT protein (L7R, G34V, S40L, D41G, D41Y, M45I, W106R, F107C and M155I).

Next, we determined the thermal stability of the remaining eighteen variants as holo-proteins (i.e. those that were expressed well as soluble proteins and were stable during purification) (Figure 3D and 4, Table S1). Nine variants showed a thermal stability close that of the WT protein (*ΔT*_m_ ≤ 2 °C; variants G3S, G3D, V9I, A29T, K32N, E36K, F107C, M155I and H162N), whereas five variants decreased the stability moderately (Δ*T*_m_ ≤ 2–5 °C; variants T16M, M45L, M45I, I51V and W106C) and four largely decrease the stability (Δ*T*_m_ ≤ 5–10 °C; Y20N, D41G, D41Y and W106R) (Table S1). Here, we note that the reported *T*_m_ and Δ*T*_m_ values are “apparent” values that cannot be regarded as reporting effects on thermodynamic stability since thermal unfolding is irreversible and kinetically-controlled (42). The W106R and W106C variants show different effects, highlighting the importance of both the location and the nature of the chemical change. Both substitutions are non-conservative changes at a residue in the active site with low solvent exposure and a low structural stability with strong ligand-dependent responsiveness based on HDX studies (27). However, their effects are very different, with W106R causing a much larger decrease in stability than W106C.

To end this section, the observed effects pinpoint that some variants in both the COSMIC and gnomAD databases decrease solubility and conformational stability of NQO1, and overall the two groups do so to similar extents (Figure 3); this observation was also seen in the computational predictions (Figure 1).

### 3.4. FAD binding and the stability of the FAD binding site

Sixteen out of the eighteen variants that yielded good levels of soluble proteins were prepared as apo-proteins to determine their affinity for FAD by titrations monitored by tryptophan fluorescence (Figure 5A, 6 and S4, Table S2). The D41Y and D41G variants were too unstable to obtain apo-proteins in sufficient amounts and quality.

**Figure 5.**
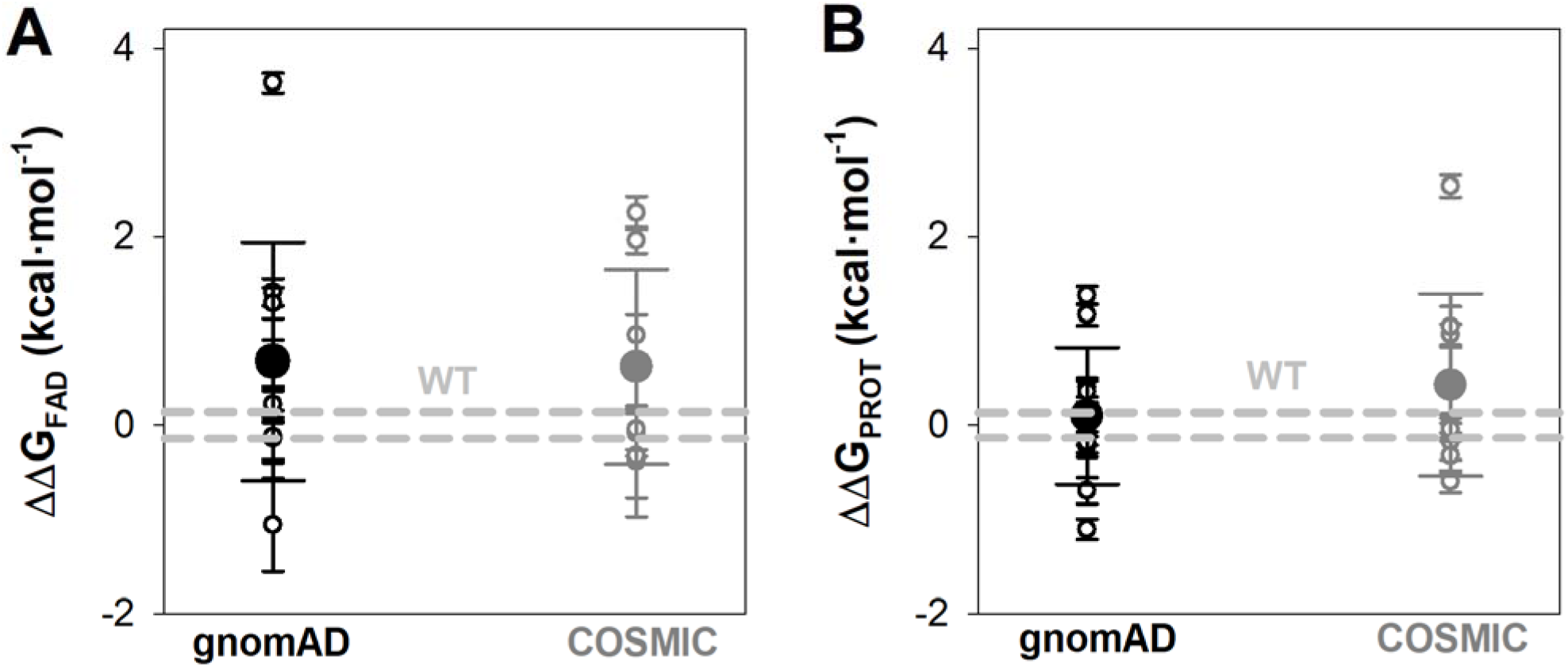
Overall effects of gnomAD and COSMIC mutations on the FAD binding affinity and the local stability of the FAD binding site (FBS). A) Changes in apparent FAD binding free energies (ΔΔG_FAD_) calculated from the difference of binding affinity between WT and a given variant from titrations. Errors are those from linear propagation; B) Local stability of the TCS (next to the FBS) from proteolysis kinetics. Changes in local stability (ΔΔG_PROT_) calculated from the difference of the second-order rate constant between WT and a given variant. Errors are those from linear propagation; For reference, the values corresponding to WT NQO1 are shown in light grey. Variants are grouped in the gnomAD and COSMIC sets.

**Figure 6.**
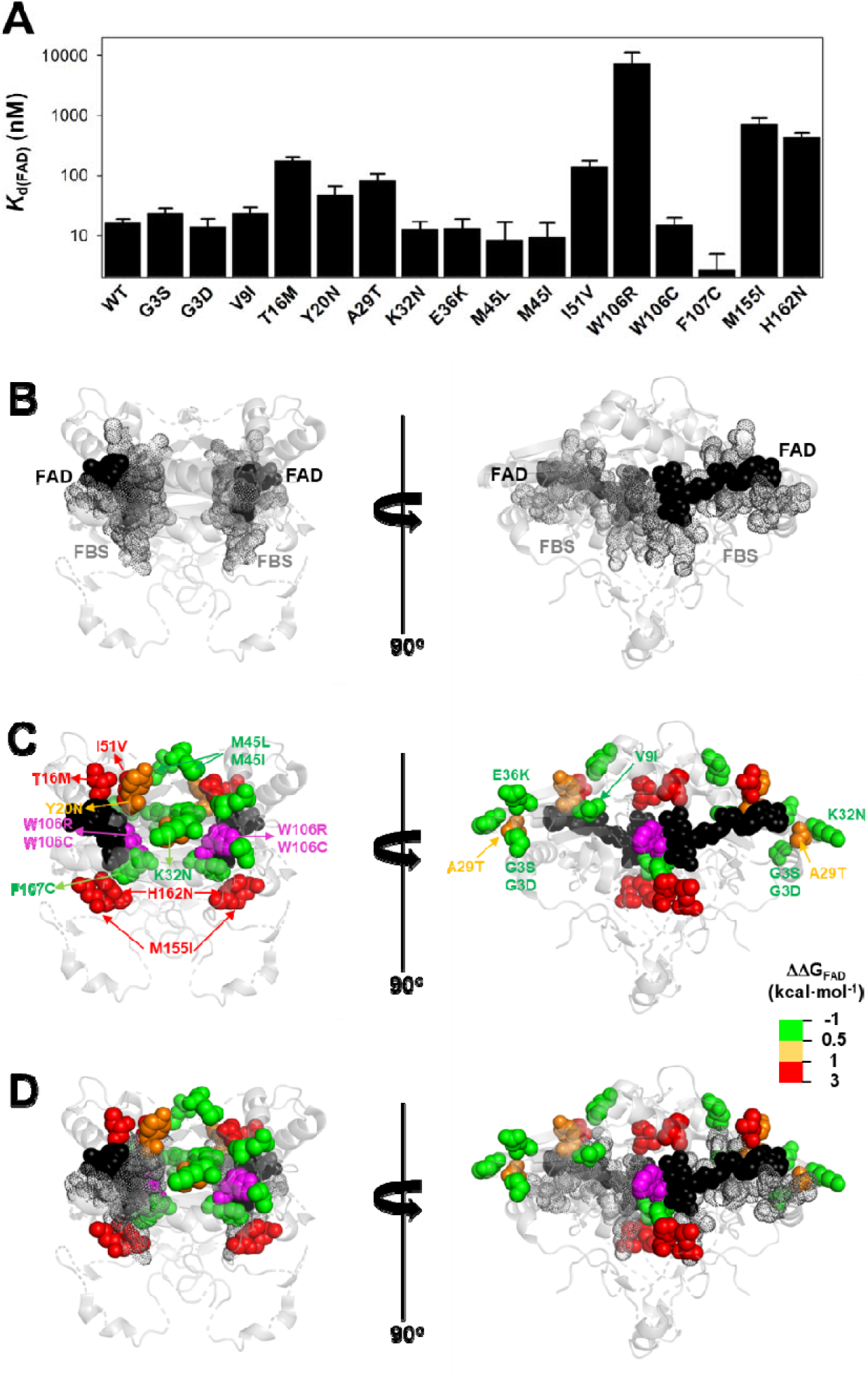
Variant effects on the FAD binding affinity. A) FAD binding affinity of each variant was determined by titrations of apo-proteins. At least two independent experiments were carried out for each variant and fitted using a single-type-of-independent binding sites to obtain *K*_d_ values (note the logarithmic scale of the y-axis). Errors are those fittings. These *K*_d_ values were used to calculate the binding free energy difference (ΔΔG_FAD_) between a given variant and the WT protein (note that a positive value indicates lower affinity). B-D) Structural location of the FAD (black spheres) and the FBS (grey dots) (panel B) and mutated residues (colour scale according to their destabilizing effect on FAD binding) (panel C). The residue W106 is highlighted in magenta due to the widely different effects of the W106R/W106C substitutions. Panel D shows an overlay of panels B and C. Note that two views rotated 90° are shown. The structural model used for display was PDB code 2F1O (54).

The G3S, G3D, V9I, K32N, E36K, M45L, M45I, W106C and F107C variants showed less than a 0.5 kcal·mol^−1^ increase in FAD binding free energy (corresponding to less than a 2.5-fold increase in *K*_d_). The Y20N and A29T variants showed moderate decrease in binding affinity (between 3-5-fold higher *K*_d_; thus, a change in FAD binding free energy of 0.5-1.0 kcal·mol^−1^). Five variants (T16M, I51V, W106R, M155I and H162N) markedly decreased the binding affinity for FAD (10-500-fold increase in *K*_d_; between 1 and 3.7 kcal·mol^−1^ decrease in binding free energy). Inspection of a structural model of NQO1 allows us to rationalize the effect of these substitutions due to their proximity to the bound FAD, with some exceptions. For instance, the W106R, W106C and F107C substitutions are in proximity to the FAD molecule, but have widely different effects (from ca. 500-fold lower affinity in W106R, to WT-like affinity for W106C and even *higher* affinity than WT for F107C). These results show that NQO1 responds to natural variations very differently even at the same site (i.e. two highly non-conservative variants at the site W106).

Mutational effects on FAD binding affinity (ΔΔG_FAD_) and proteolysis rates (ΔΔG_PROT_) with thermolysin have previously been shown to correlate well (24,25,43). The site for initial cleavage by thermolysin (TCS) is generally between Ser72–Val73, close to the FAD binding site (28). All the variants investigated showed proteolysis patterns that were similar to that of WT NQO1 (Figure S5). The previously observed between ΔΔG_FAD_ and ΔΔG_PROT_ holds for the 16 variants for which both FAD binding affinity and protease sensitivity can be compared (Figure S6). The T16M, I51V, W106C and M155I mutations destabilized locally the TCS by 1–2.5 kcal·mol^−1^ and the residues affected by these substitutions are in general close to the TCS (with the exception of M155I, the most destabilizing mutation) (Figure 7). Again, the results for the W106R/C variants were very different: both affected the local stability by ~ 1 kcal·mol^−1^, but with opposite signs (Table S2).

**Figure 7.**
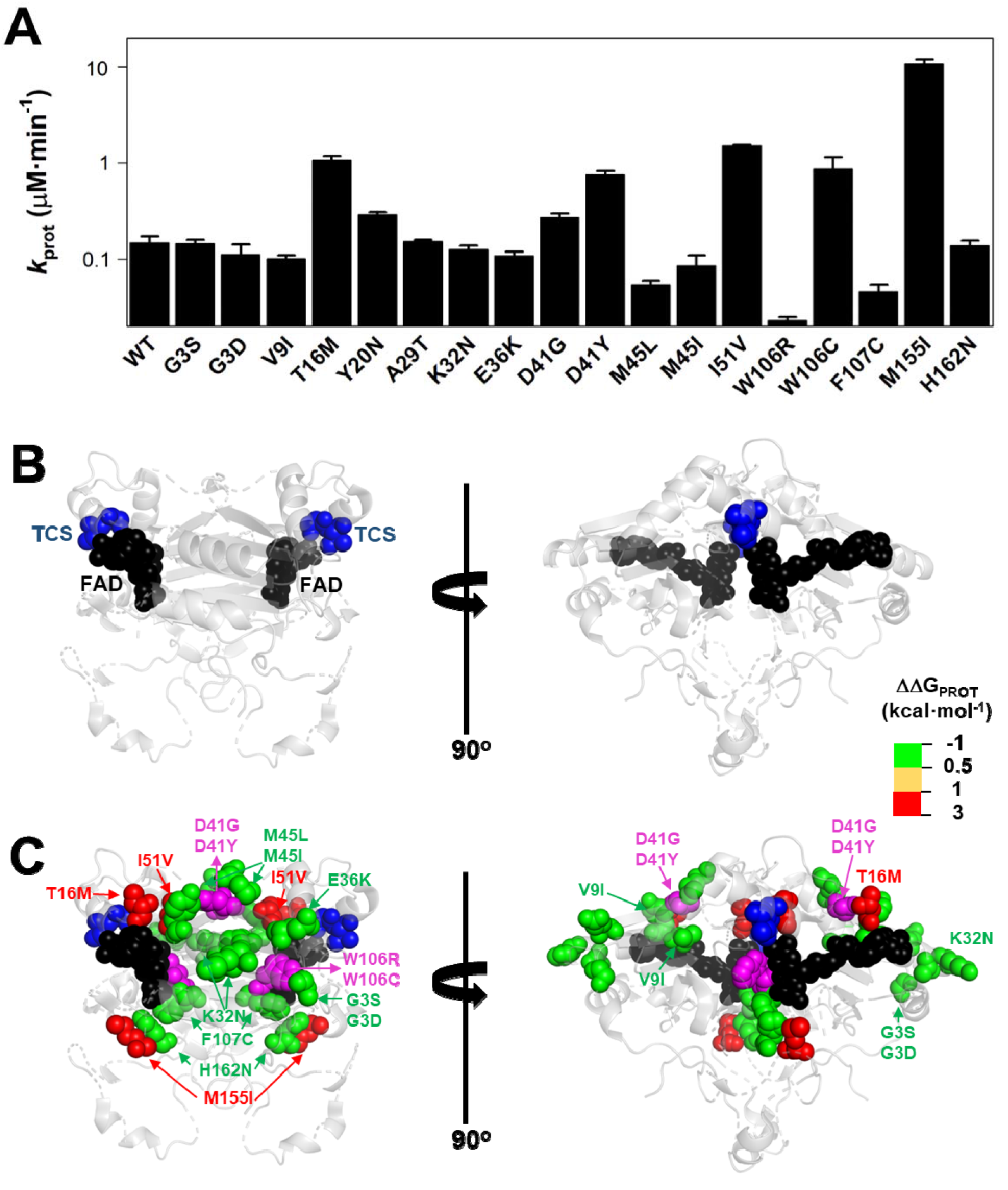
Variant effects on the local stability of the FBS from proteolysis. A) Second-order rate constants for proteolysis of NQO1 variants. Figure S5C. Errors are those fittings. These *k*_PROT_ values were used to calculate the TCS local energy free difference (ΔΔG_PROT_) between a given variant and the WT protein (note that a positive value indicates lower affinity). B-D) Structural location of the FAD (black spheres) and the TCS (blue spheres) (panel B). In C, mutated residues (color scale according to their destabilizing effect on FAD binding) (panel C) are overlayed with TCS and FAD. The residues D41 and W106 are highlighted in magenta due to the widely different effects of the D41G/D41Y and W106R/W106C substitutions. Note that two views rotated 90° are shown. The structural model used for display was PDB code 2F1O (54).

Overall, the negative impact on FAD binding affinity and the stability of the FAD binding site in the holo-state was similar between variants from COSMIC and gnomAD sets (Figure 5).

### 3.5. Comparison of experimental analyses and computational predictions

We then proceeded to compare the experimental data to each other and to computational predictions. To ease comparison between the calculated ΔΔG values and thermal melting measurements, we first estimated ΔΔG_melting_ from the Δ*T*_m_ values using an empirical relationship (44), again noting that these are not strictly thermodynamic values as unfolding was not reversible. We first compared the experimental values of ΔΔG_melting_ to the levels of soluble protein (S values, Figure 8A). We found that unstable variants (those with ΔΔG_melting_ > 2 kcal·mol^−1^ or not amenable for purification, not-determined or ND) mostly showed level of S close to zero (< 5) except for L7P. Stable variants (here defined as ΔΔG_melting_ < 2 kcal·mol^−1^) instead showed a wide range of S values (76±85%; mean ± s.d.).

**Figure 8.**
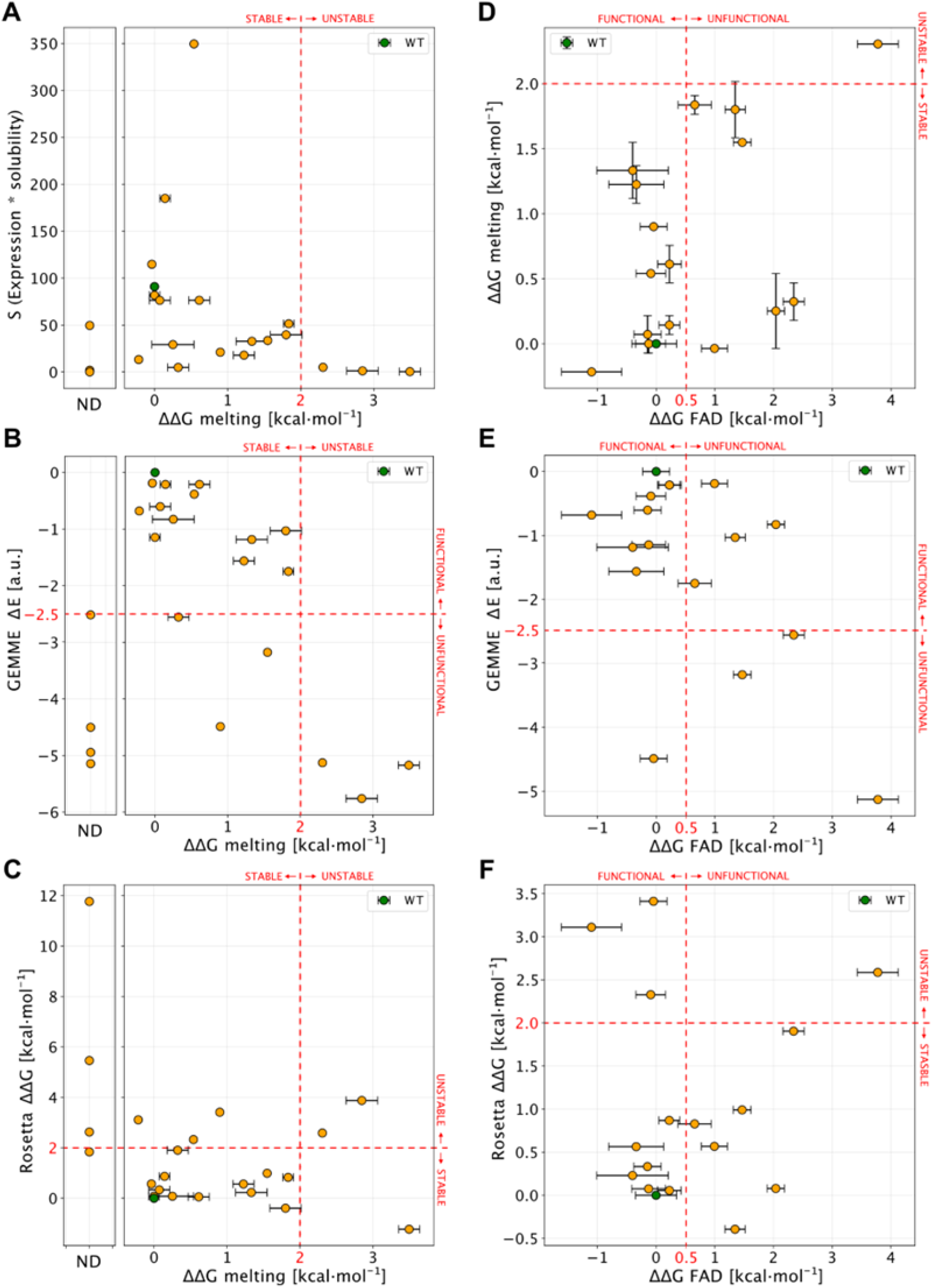
Comparison of experimental results with computational scores. A: Scatter plot of ΔΔG_melting_ (x-axis) and S values (expression * solubility). Not-determined variants (ND) in thermal stability experiment are reported in a separate plot. B and C show a comparison between ΔΔG_melting_ and computational predictors. D: Correlation between ΔΔG_melting_ and ΔΔG_FAD_ for NQO1 variants detected in both the experiments. E and F: Comparison of ΔΔG_FAD_ with the computational results. Red lines, if present, show the boundary of experimental and computational classes for each comparison. In each panel errors are reported as a black bar on every single variant, if present.

We next compared ΔΔG_melting_ with the computational scores (Figure 8B-C). Overall, we found a good agreement for most of the unstable and Not-Determined (ND) variants, which showed ΔΔG > 2.0 kcal·mol^−1^ and evolutionary distance, ΔE < −3 indicating predicted loss of stability and function. The only exception was D41Y which displayed a stabilizing behaviour in Rosetta ΔΔG predictions. Experimentally stable variants (ΔΔG_melting_ < 2 kcal·mol^−1^) displayed low evolutionary distances (ΔE > −2.5 kcal·mol^−1^) except for W106C and T16M. This observation for T16M supports the notion that detrimental effects on protein function are not connected to thermodynamic destabilization for this variant (25).

We then compared ΔΔG_FAD_ with the other experimental and computational observables (Figure 8D-F). For most (15 out of 16) of the variants where ΔΔG_FAD_ could be measured, the ΔΔG_melting_ was < 2 kcal·mol^−1^, indicating stable variants which was also confirmed by Rosetta ΔΔG calculations (13 out of 16 substitutions). Seven of the fifteen variants showed a ΔΔG_FAD_ > 0.5 kcal·mol^−1^ indicating loss of function. Of these, three variants were captured by evolutionary conservation analysis (ΔE < −2.5 kcal·mol^−1^).

To summarize, for 14 out of 22 tested variants (G3S L7P L7D V9I T16M Y20N K32N G34V E36K D41G M45L M45I W106R, M155I) the predictions from the computational protocols match the experimental results, reinforcing the evidences of the effects of the variants on the protein stability and function.

While the results are overall encouraging, there remains differences between computation and experiments for the effects of some mutations (eight out of twenty-two; G3D, A29T, S40L, D41Y, I51V, W106C, F107C and H162N). For five of these (G3D, S40L, D41Y, W106C and F107C) it appears that there is a difference between the stability prediction by Rosetta and experiments (noting again that the latter are not equilibrium measurements). For the partially exposed S40L and D41Y the reason for the discrepancy is perhaps related to specific interactions made by these two residues whose effects are not captured by the Rosetta calculations. Both W106C and F107C involve substituting aromatic residues with a cysteine, suggesting problems with evaluating such substitutions. In other two cases (G3D, A29T and H162N) the GEMME scores did not capture properly the variant effects, possibly because some specific interactions in human NQO1 may not be present in other homologs of NQO1 and thus, not captured by the evolutionary analysis. Lastly, for I51V the behaviour is opposite from both computational predictions.

## 4. Discussion

With advances in sequencing technologies, we are uncovering the large genetic variability in the *human genome*. To exploit the availability of this information at the clinical level, we must be able to establish genotype-phenotype correlations accurately and at a large-scale. Although detailed characterization of mutational effects is obviously useful, it is difficult to perform this at such scale (many genes, many variants). However, we may use experimental characterization on a more modest scale to test the performance of current predictive tools in the hope that we can improve them. In this work, we have carried out such an exercise for the human NQO1 protein. The rationale for selecting this system is three-fold: i) human NQO1 is a multifunctional protein and mutational effects may affect these functions through complex mechanisms (30,45). Therefore, contrasting experimental characterization and computational predictions can be challenging for current predictive tools and may help to improve them; ii) Altered NQO1 functionality is associated with increased risk of developing cancer and neurological disorders (13). Indeed, the presence of a highly deleterious polymorphic variant in NQO1 is associated with increased cancer risk and affects multiple functions through allosteric effects (45,46); iii) Over a hundred of missense variants in human NQO1 have been found in human population (i.e. the gnomAD database) or in cancer cell lines as somatic mutations (i.e. the COSMIC database). However, the impact of these mutations on NQO1 multifunctionality and their potential role in cancer development are largely unexplored.

Theoretical advances and new methodologies in the fields of sequence evolution and structure predictions allow us to perform large-scale *in silico* mutagenesis studies on target proteins. Although the current state-of-art algorithms are often (but not always (47)) less accurate at predicting pathogenicity compared to detailed experimental testing, they provide a fast and effective way to predict LoF and sometimes to generate mechanistic hypotheses regarding which properties a variant might affect (8,48).

Here, we first performed *in silico* saturation mutagenesis of WT NQO1, predicting the changes in thermodynamic stability (ΔΔG) and evolutionary conservation (ΔE) for 5187 variants. We combined the two scores to perform a global analysis on how the NQO1 function may be perturbed. Approximately 44% of variants are predicted to cause loss of function, with 45% of these drastically affecting the protein stability. This analysis enabled us to obtain an overview on the possible biologically relevant positions and variants. Indeed, although we know that the ability of computational tools to assign biological functions and predict overall pathogenicity is rapidly increasing, we are still at a point where computational methods may not predict LoF perfectly, and often do not shed much light on the mechanisms of action. This might in particular hold for proteins like NQO1 where multiple biological functions are present, and where some of which may differ between orthologues.

We then used the information provided from the *in silico* saturation mutagenesis to select 22 naturally-occurring variants with a diverse range of predicted effects on protein stability and function to be experimentally tested. We selected nine mutations found in COSMIC and thirteen from gnomAD. Of these variants, 36% severely affected protein foldability and solubility (upon expression in *E.coli*) or reduced conformational stability (at least a 5 °C decrease in *T*_m_). A quarter of the variants had severely affected FAD binding (a 5-fold decrease in affinity, i.e. a 1 kcal·mol^−1^ of binding free energy penalization). For 64% of the variants, experimental characterization and computational predictions agreed in the variant effects on protein stability and function, whereas the remaining 36% of the mutations might be explained by limitations known for the tools used in the prediction process. Although, at this point, this level of agreement is reasonable, it also pinpoints the necessity of improving these predictive tools.

COSMIC mutations are in general somatic (actually, 86% of the COSMIC mutations of NQO1 are labelled as *confirmed* in this database; accessed by 17^th^ August 2022) and likely come from samples that underwent many mutational events in different genes. Thus, the identification of a mutation in the COSMIC database does not imply that this mutation is a driver mutation (here we may define a driver mutation as a mutation with the ability to drive tumorigenesis and confer selective advantages in a tumor cell and a somatic tissue; (49)). Mutations in the gnomAD database belong to heterogeneous groups (many different sequencing projects, some of them case-control studies), and likely reflect genetic variability in the *germline* and in general presence or absence in gnomAD is not sufficient to assign a label as pathogenic or benign. When we examine the NQO1 variants investigated in this work found in the gnomAD database (v.2.1.), allele frequencies are overall comparable in *control* vs. *all* samples (Table S3). This suggests that there is no strong bias towards *case* samples, and thus the allele frequencies in gnomAD may represent well their presence in a *healthy* population. The presence of these mutations in the germline may predispose somatic cells towards a new mutational event in the WT allele (as occurs in familial cancer cases; (49)), thus largely decreasing the NQO1 activity and function.

Our combined experimental and computational analyses provide information on the potential LoF character and the mechanisms by which the variants may exert their effects (protein stability and/or function). Due to its role in the antioxidant defense and stabilization of oncosuppressor proteins, it is likely that NQO1 play a role in cancer development. Homozygous NQO1 knock-out mice revealed cancer-associated phenotypic traits when exposed to chemical or radiological insults (50–53). Thus, the presence of LoF variants in NQO1 and increased cancer risk may resemble a recessive inheritance (46). The p.P187S polymorphism (with an allele frequency of ~0.25, Table S3) dramatically decreases the intracellular stability of NQO1 thus preventing its interaction with oncosuppressors, reducing enzyme activity and affecting almost the entire structure of NQO1 (5). Noteworthy, it only associates with cancer in homozygotes (46). Due to the low frequency of most gnomAD NQO1 variants, their presence would be rare even in compound heterozygotes. In fact, 98% of the homozygous samples containing NQO1 missense variations correspond to homozygotes for P187S. However, an additional (*somatic*) mutational event in a WT/P187S genetic background (about 25% of human population) may substantially enhance the LOF phenotype in this common genetic background.

We present a test of predictive tools against the experimental characterization of large set of naturally-occurring mutations on NQO1 stability and function. Further steps will be taken to provide a wider perspective on the multifunctionality of NQO1 (i.e. intracellular degradation and stability, high-resolution structural stability in different ligation states, enzyme function and cooperativity, interaction with protein partners, allosteric communication of mutational effects) and the relationships between the genetic diversity of NQO1 in human population and its link with individual propensity towards disease development.

## Data availability statement

The original contributions presented in the study are included in the article/Supplementary Material, further inquiries can be directed to the corresponding author.

## Author contributions

A.L.P. conceived the project and supervised the experimental work; J.L.P-G. and K. J-T. carried out expression, purification and characterization of proteins; M.C. carried out computational analysis; E.S. contributed to selection of variants and contributed with reagents; K.L-L. supervised the computational work; E.S., K.L-L. and A.L.P. received funding; M.C., K.L-L. and ALP wrote the original draft; All authors contributed to and approved the final version of the manuscript.

## References

1. Arnedo-Pac C, Lopez-Bigas N, Muiños F. (2022). Predicting disease variants using biodiversity and machine learning. Nat. Biotech. 40, 27–8.

2. Høie MH, Cagiada M, Beck Frederiksen AH, Stein A, Lindorff-Larsen K. (2022). Predicting and interpreting large-scale mutagenesis data using analyses of protein stability and conservation. Cell Rep. 38, 110207. doi: 10.1016/j.celrep.2021.110207.

3. Katsonis P, Wilhelm K, Williams A, Lichtarge O. (2022). Genome interpretation using in silico predictors of variant impact. Hum. Genet. doi: 10.1007/s00439-022-02457-6.

4. McInnes G, Sharo AG, Koleske ML, Brown JEH, Norstad M, Adhikari AN, et al. (2021). Opportunities and challenges for the computational interpretation of rare variation in clinically important genes. Am. J. Hum. Genet. 108, 535–48.

5. Pacheco-Garcia JL, Anoz-Carbonell E, Vankova P, Kannan A, Palomino-Morales R, Mesa-Torres N, et al. (2021). Structural basis of the pleiotropic and specific phenotypic consequences of missense mutations in the multifunctional NAD(P)H:quinone oxidoreductase 1 and their pharmacological rescue. Redox Biol. 46:102112. doi: 10.1016/j.redox.2021.102112.2021.

6. Xu Q, Tang Q, Katsonis P, Lichtarge O, Jones D, Bovo S, et al. (2017). Benchmarking predictions of allostery in liver pyruvate kinase in CAGI4. Hum. Mutat. 38, 1123–31.

7. Abildgaard AB, Stein A, Nielsen SV, Schultz-Knudsen K, Papaleo E, Shrikhande A, et al. (2019). Computational and cellular studies reveal structural destabilization and degradation of MLH1 variants in Lynch syndrome. Elife. 8, e49138. doi: 10.7554/eLife.49138.

8. Cagiada M, Johansson KE, Valanciute A, Nielsen S v, Hartmann-Petersen R, Yang JJ, et al. (2021). Understanding the Origins of Loss of Protein Function by Analyzing the Effects of Thousands of Variants on Activity and Abundance. Mol. Biol. Evol. 38, 3235–46.

9. Luo S, Kang SS, Wang ZH, Liu X, Day JX, Wu Z, et al. (2019). Akt phosphorylates NQO1 and triggers its degradation, abolishing its antioxidative activities in Parkinson’s disease. J. Neurosci. 39, 7291–305.

10. Beaver SK, Mesa-Torres N, Pey AL, Timson DJ. (2019). NQO1: A target for the treatment of cancer and neurological diseases, and a model to understand loss of function disease mechanisms. Biochim. Biophys. Acta Proteins Proteom. 1867, 663–76.

11. Ross D, Siegel D. (2018) NQO1 in protection against oxidative stress. Curr. Opin Toxicol. 7, 67–72.

12. Anoz-Carbonell E, Timson DJ, Pey AL, Medina M. (2020). The catalytic cycle of the antioxidant and cancer-associated human NQO1 enzyme: Hydride transfer, conformational dynamics and functional cooperativity. Antioxidants. 9, 772. doi: 10.3390/antiox9090772.

13. Salido E, Timson DJ, Betancor-Fernández I, Palomino-Morales R, Anoz-Carbonell E, Pacheco-García JL, et al. (2022). Targeting HIF-1α Function in Cancer through the Chaperone Action of NQO1: Implications of Genetic Diversity of NQO1. J. Pers Med. 12, 747. doi: 10.3390/jpm12050747.12,.

14. Asher G, Tsvetkov P, Kahana C, Shaul Y. (2005). A mechanism of ubiquitin-independent proteasomal degradation of the tumor suppressors p53 and p73. Genes Dev. 19, 316–21.

15. Ben-Nissan G, Sharon M. (2014). Regulating the 20S proteasome ubiquitin-independent degradation pathway. Biomolecules. 4, 862–84. doi: 10.3390/biom4030862.

16. di Francesco A, di Germanio C, Panda AC, Huynh P, Peaden R, Navas-Enamorado I, et al. (2016). Novel RNA-binding activity of NQO1 promotes SERPINA1 mRNA translation. Free Radic. Biol. Med. 99, 225–33.

17. Oh ET, Kim JW, Kim JM, Kim SJ, Lee JS, Hong SS, et al. (2016). NQO1 inhibits proteasome-mediated degradation of HIF-1α. Nat. Commun. 7:13593. doi: 10.1038/ncomms13593.

18. Martínez-Limón A, Calloni G, Ernst R, Vabulas RM. (2020). Flavin dependency undermines proteome stability, lipid metabolism and cellular proliferation during vitamin B2 deficiency. Cell. Death. Dis. 11, 725. doi: 10.1038/s41419-020-02929-5.

19. Martínez-Limón A, Alriquet M, Lang WH, Calloni G, Wittig I, Vabulas RM. (2016). Recognition of enzymes lacking bound cofactor by Protein quality control. Proc. Natl. Acad. Sci. U S A. 113:12156–61.

20. Siegel D, Bersie S, Harris P, di Francesco A, Armstrong M, Reisdorph N, et al. (2021). A redox-mediated conformational change in NQO1 controls binding to microtubules and α-tubulin acetylation. Redox Biol. 39, 101840. doi: 10.1016/j.redox.2020.101840.

21. Faig M, Bianchet MA, Talalay P, Chen S, Winski S, Ross D, et al. (2000). Structures of recombinant human and mouse NAD(P)H:quinone oxidoreductases: Species comparison and structural changes with substrate binding and release. Proc. Natl. Acad. Sci. U S A. 97, 3177–3182.

22. Li R, Bianchet MA, Talalayt P, Amzel LM. (1995). The three-dimensional structure of NAD(P)H:quinone reductase, a flavoprotein involved in cancer chemoprotection and chemotherapy: Mechanism of the two-electron reduction. Proc. Natl. Acad. Sci. U S A. 92, 8846–8850.

23. Lienhart WD, Gudipati V, Uhl MK, Binter A, Pulido SA, Saf R, et al. (2014). Collapse of the native structure caused by a single amino acid exchange in human NAD(P)H:Quinone oxidoreductase. FEBS J. 281, 4691–704.

24. Medina-Carmona E, Neira JL, Salido E, Fuchs JE, Palomino-Morales R, Timson DJ, et al. (2017). Site-to-site interdomain communication may mediate different loss-of-function mechanisms in a cancer-associated NQO1 polymorphism. Sci. Rep. 7, 44532. doi: 10.1038/srep44532.

25. Pacheco-García JL, Cano-Muñoz M, Sánchez-Ramos I, Salido E, Pey AL. (2020). Naturally-occurring rare mutations cause mild to catastrophic effects in the multifunctional and cancer-associated NQO1 protein. J. Pers. Med. 10, 1–31. doi: 10.3390/jpm10040207.

26. Pey AL. Biophysical and functional perturbation analyses at cancer-associated P187 and K240 sites of the multifunctional NADP(H):quinone oxidoreductase 1. Int. J. Biol. Macromol. 2018 Oct 15;118:1912–23.

27. Vankova P, Salido E, Timson DJ, Man P, Pey AL. (2019). A dynamic core in human NQO1 controls the functional and stability effects of ligand binding and their communication across the enzyme dimer. Biomolecules. 9, 728. doi: 10.3390/biom9110728.

28. Medina-Carmona E, Palomino-Morales RJ, Fuchs JE, Padín-Gonzalez E, Mesa-Torres N, Salido E, et al. (2016). Conformational dynamics is key to understanding loss-of-function of NQO1 cancer-associated polymorphisms and its correction by pharmacological ligands. Sci. Rep. 2016 Apr 22;6(1):20331.

29. Medina-Carmona E, Betancor-Fernández I, Santos J, Mesa-Torres N, Grottelli S, Batlle C, et al. (2019). Insight into the specificity and severity of pathogenic mechanisms associated with missense mutations through experimental and structural perturbation analyses. Hum. Mol. Genet. 28, 1–15.

30. Pacheco-Garcia JL, Anoz-Carbonell E, Loginov DS, Vankova P, Salido E, Man P, et al. (2022). Different phenotypic outcome due to site-specific phosphorylation in the cancer-associated NQO1 enzyme studied by phosphomimetic mutations. Arch. Biochem. Biophys. 729, 109392.

31. Medina-Carmona E, Fuchs JE, Gavira JA, Mesa-Torres N, Neira JL, Salido E, et al. (2017). Enhanced vulnerability of human proteins towards disease-associated inactivation through divergent evolution. Hum. Mol. Genet. 26, 3531–44.

32. Park C, Marqusee S. (2004). Probing the high energy states in proteins by proteolysis. J. Mol. Biol. 343, 1467–76.

33. Laine E, Karami Y, Carbone A. (2019). GEMME: A Simple and Fast Global Epistatic Model Predicting Mutational Effects. Mol. Biol. Evol. 36, 2604–2619. doi: 10.1093/molbev/msz179.

34. Steinegger M, Meier M, Mirdita M, Vöhringer H, Haunsberger SJ, Söding J. (2019). HH-suite3 for fast remote homology detection and deep protein annotation. BMC Bioinformatics. 20, 473.

35. Remmert M, Biegert A, Hauser A, Söding J. (2011). HHblits: lightning-fast iterative protein sequence searching by HMM-HMM alignment. Nat. Methods. 9, 173–5.

36. Frenz B, Lewis SM, King I, DiMaio F, Park H, Song Y. (2020). Prediction of Protein Mutational Free Energy: Benchmark and Sampling Improvements Increase Classification Accuracy. Front. Bioeng. Biotechnol. 8, 558247. doi: 10.3389/fbioe.2020.558247.

37. Park H, Bradley P, Greisen P, Liu Y, Mulligan VK, Kim DE, et al. (2016). Simultaneous Optimization of Biomolecular Energy Functions on Features from Small Molecules and Macromolecules. J. Chem. Theory Comput. 12, 6201–12.

38. Kabsch W, Sander C. (1983). Dictionary of protein secondary structure: Pattern recognition of hydrogen-bonded and geometrical features. Biopolymers. 22, 2577–637.

39. Naganathan AN. (2019). Modulation of allosteric coupling by mutations: from protein dynamics and packing to altered native ensembles and function. Curr. Opin. Struct. Biol. 54, 1–9.

40. Luque I, Freire E. (2000). Structural stability of binding sites: consequences for binding affinity and allosteric effects. Proteins. Suppl 4, 63–71.

41. Luque I, Leavitt SA, Freire E. (2002). The linkage between protein folding and functional cooperativity: Two sides of the same coin?. Annu. Rev. Biophys. Biomol. Struct. 31, 235–56.

42. Pey AL, Megarity CF, Timson DJ. (2004). FAD binding overcomes defects in activity and stability displayed by cancer-associated variants of human NQO1. Biochim. Biophys. Acta Mol. Basis Dis. 1842, 2163–73.

43. Medina-Carmona E, Rizzuti B, Martín-Escolano R, Pacheco-García JL, Mesa-Torres N, Neira JL, et al. (2019). Phosphorylation compromises FAD binding and intracellular stability of wild-type and cancer-associated NQO1: Insights into flavo-proteome stability. Int. J. Biol. Macromol. 125, 1275–88.

44. Watson MD, Monroe J, Raleigh DP. (2018). Size-Dependent Relationships between Protein Stability and Thermal Unfolding Temperature Have Important Implications for Analysis of Protein Energetics and High-Throughput Assays of Protein-Ligand Interactions. J. Phys. Chem. B. 122, 5278–85.

45. Pacheco-Garcia JL, Loginov DS, Anoz-Carbonell E, Vankova P, Palomino-Morales R, Salido E, et al. (2022). Allosteric Communication in the Multifunctional and Redox NQO1 Protein Studied by Cavity-Making Mutations. Antioxidants. 11, 1110. doi: 10.3390/antiox11061110.

46. Lajin B, Alachkar A. (2013). The NQO1 polymorphism C609T (Pro187Ser) and cancer susceptibility: a comprehensive meta-analysis. Br. J. Cancer. 109, 1325–37.

47. Frazer J, Notin P, Dias M, Gomez A, Min JK, Brock K, et al. (2021). Disease variant prediction with deep generative models of evolutionary data. Nature. 599, 91–5.

48. Stein A, Fowler DM, Hartmann-Petersen R, Lindorff-Larsen K. (2019). Biophysical and Mechanistic Models for Disease-Causing Protein Variants. Trends Biochem. Sci. 44, 575–88.

49. Martínez-Jiménez F, Muiños F, Sentís I, Deu-Pons J, Reyes-Salazar I, Arnedo-Pac C, et al. (2020). A compendium of mutational cancer driver genes. Nat. Rev. Cancer. 20, 555–72.

50. Long DJ, Waikel RL, Wang XJ, Perlaky L, Roop DR, Jaiswal AK. (2000). NAD(P)H:quinone oxidoreductase 1 deficiency increases susceptibility to benzo(a)pyrene-induced mouse skin carcinogenesis. Cancer Res. 60, 5913–5.

51. Iskander K, Gaikwad A, Paquet M, Long DJ, Brayton C, Barrios R, et al. (2005). Lower induction of p53 and decreased apoptosis in NQO1-null mice lead to increased sensitivity to chemical-induced skin carcinogenesis. Cancer Res. 65, 2054–8.

52. Iskander K, Barrios RJ, Jaiswal AK. (2008). Disruption of NAD(P)H:quinone oxidoreductase 1 gene in mice leads to radiation-induced myeloproliferative disease. Cancer Res. 68, 7915–22.

53. Radjendirane V, Joseph P, Lee YH, Kimura S, Klein-Szanto AJ, Gonzalez FJ, et al. (1998). Disruption of the DT diaphorase (NQO1) gene in mice leads to increased menadione toxicity. J. Biol. Chem. 273, 7382–9.

54. Asher G, Dym O, Tsvetkov P, Adler J, Shaul Y. (2006). The crystal structure of NAD(P)H quinone oxidoreductase 1 in complex with its potent inhibitor dicoumarol. Biochemistry. 45, 6372–8.

