## Supplementary Material for "Effect of naturally-occurring mutations on the stability and function of cancer-associated NQO1: comparison of experiments and computation"

**Supplementary Table 1. Variant effects on the solubility and thermal stability of NQO1.** Expression analyses of NQO1 was carried out in *E.coli* at 37^o^C and the total expression levels (T), and the fraction of soluble protein (S/T ratio) were determined by western-blot (mean ± s.d. from three independent experiments). The % of soluble protein (S) was thus determined as the T x S/T product. *T*_m_ values were determined upon purification of NQO1 proteins by thermal denaturation (mean±s.d. from at least three replicates). N.Det. not determined due to very low expression levels as soluble protein.

| **Variant** | **T**  **(% vs.WT)** | **S/T ratio** | **S**  **(%)** | ***T*_m_ (^o^C)** |
| --- | --- | --- | --- | --- |
| WT | 100 | 0.91±0.13 | 91±13 | 51.8±0.5 |
| G3S | 185±26 | 1.00±0.07 | 185±27 | 51.4±0.6* |
| G3D | 454±103 | 0.77±0.10 | 350±103 | 50.3±0.5* |
| L7P | 620±97 | 0.08±0.04 | 50±10 | N.Det. |
| L7R | 9±10 | 0.11±0.03 | ~ 1 | N.Det. |
| V9I | 156±48 | 0.49±0.37 | 76±60 | 50.1±0.3* |
| T16M | 96±10 | 0.35±0.12 | 34±16 | 47.5±0.5* |
| Y20N | 97±10 | 0.53±0.08 | 51±13 | 46.7±0.4* |
| A29T | 129±26 | 0.89±0.11 | 115±28 | 51.9±0.5* |
| K32N | 134±7 | 0.52±0.17 | 76±18 | 51.6±0.3 |
| G34V | 6±1 | 0.32±0.11 | ~ 2 | N.Det. |
| E36K | 94±11 | 0.87±0.09 | 82±14 | 51.8±0.6 |
| S40L | 2±1 | 0.04±0.03 | > 1 | N.Det. |
| D41G | 7±2 | 0.18±0.23 | ~ 1 | 43.9±0.2 |
| D41Y | 2±1 | 0.21±0.14 | ~ 1 | 42.1±0.3 |
| M45L | 52±41 | 0.63±0.07 | 33±42 | 48.1±0.2 |
| M45I | 46±29 | 0.39±0.26 | 18±39 | 48.4±0.3 |
| I51V | 64±21 | 0.62±0.13 | 40±25 | 46.8±0.2 |
| W106R | 24±16 | 0.21±0.07 | ~ 5 | 45.4±0.5 |
| W106C | 66±45 | 0.32±0.08 | 21±45 | 49.3±0.5 |
| F107C | 95±64 | 0.14±0.10 | 13±65 | 52.4±0.5 |
| M155I | 35±33 | 0.14±0.11 | ~ 5 | 50.9±0.3 |
| H162N | 65±71 | 0.45±0.38 | 29±80 | 51.1±0.9 |

* From (1).

**Supplementary Table 2. Variant effects on FAD binding.** The apparent dissociation constants (*K*_d FAD_) were determined by titrations of apo-NQO1 variants with FAD. Data are best-fit parameters from at least two independent titrations. The local stability of the TCS (close to the FAD binding site) was determined by proteolysis with thermolysin (*k*_prot_). *k*_prot_ is the second-order rate constant for proteolysis obtained from the linear dependence of the apparent first-order rate constant on protease concentration.

| **Variant** | ***K*_d_ _FAD_ (nM)** | ***k*_prot_** **(µM·min^-1^)** |
| --- | --- | --- |
| WT | 16.1±2.7 | 0.149±0.024 |
| G3S | 23.2±5.4 | 0.146 ± 0.013 |
| G3D | 13.9±5.2 | 0.111 ± 0.032 |
| L7P | N.Det. | N.Det. |
| L7R | N.Det. | N.Det. |
| V9I | 23.3±6.5 | 0.101 ± 0.009 |
| T16M | 174±31 | 1.075 ± 0.115 |
| Y20N | 46.8±20.0 | 0.292 ± 0.017 |
| A29T | 80.8±25.7 | 0.153 ± 0.005 |
| K32N | 12.7±4.3 | 0.126±0.013 |
| G34V | N.Det. | N.Det. |
| E36K | 13.1±5.7 | 0.109±0.011 |
| S40L | N.Det. | N.Det. |
| D41G | N.Det. | 0.271 ± 0.029 |
| D41Y | N.Det. | 0.754 ± 0.072 |
| M45L | 8.4±8.2 | 0.054 ± 0.006 |
| M45I | 9.3±6.9 | 0.086 ± 0.025 |
| I51V | 143±32 | 1.521 ± 0.032 |
| W106R | 7400±4000* | 0.023±0.002 |
| W106C | 15±5* | 0.864±0.285 |
| F107C | 2.7±2.2* | 0.046±0.008 |
| M155I | 722±173* | 10.75 ± 1.26 |
| H162N | 441±74* | 0.138±0.018 |

* Unpublished work.

**Supplementary Table 3. Allelic frequency of NQO1 mutations found in the gnomAD database and experimentally analysed in this work.** Frequencies are reported from the gnomAD v.2.1. using all or control samples. For sake of comparison, data on the common P187S polymorphism is also added.

| **Variation** | **Allelic frequency** | |
| --- | --- | --- |
|  | **All (141.456) samples** | **Control (60.146) samples** |
| **G3S** | 1.21·10^-5^ | 1.86·10^-5^ |
| **L7R** | 7.80·10^-5^ | 5.49·10^-5^ |
| **V9I** | 3.90·10^-5^ | 4.16·10^-5^ |
| **T16M** | 2.83·10^-5^ | 4.64·10^-4^ |
| **Y20N** | 2.12·10^-5^ | 1.83·10^-5^ |
| **K32N** | 7.96·10^-5^ | 4.57·10^-5^ |
| **G34V** | 3.98·10^-6^ | 9.14·10^-6^ |
| **E36K** | 6.37·10^-5^ | 3.33·10^-5^ |
| **S40L** | 1.99·10^-5^ | 9.14·10^-6^ |
| **D41G** | 3.98·10^-6^ | 9.14·10^-6^ |
| **I51V** | 6.37·10^-5^ | 3.33·10^-5^ |
| **W106R** | 2.79·10^-5^ | 4.57·10^-5^ |
| **F107C** | 1.77·10^-5^ | 0 |
| **P187S** | 2.47·10^-1^ | 2.55·10^-1^ |

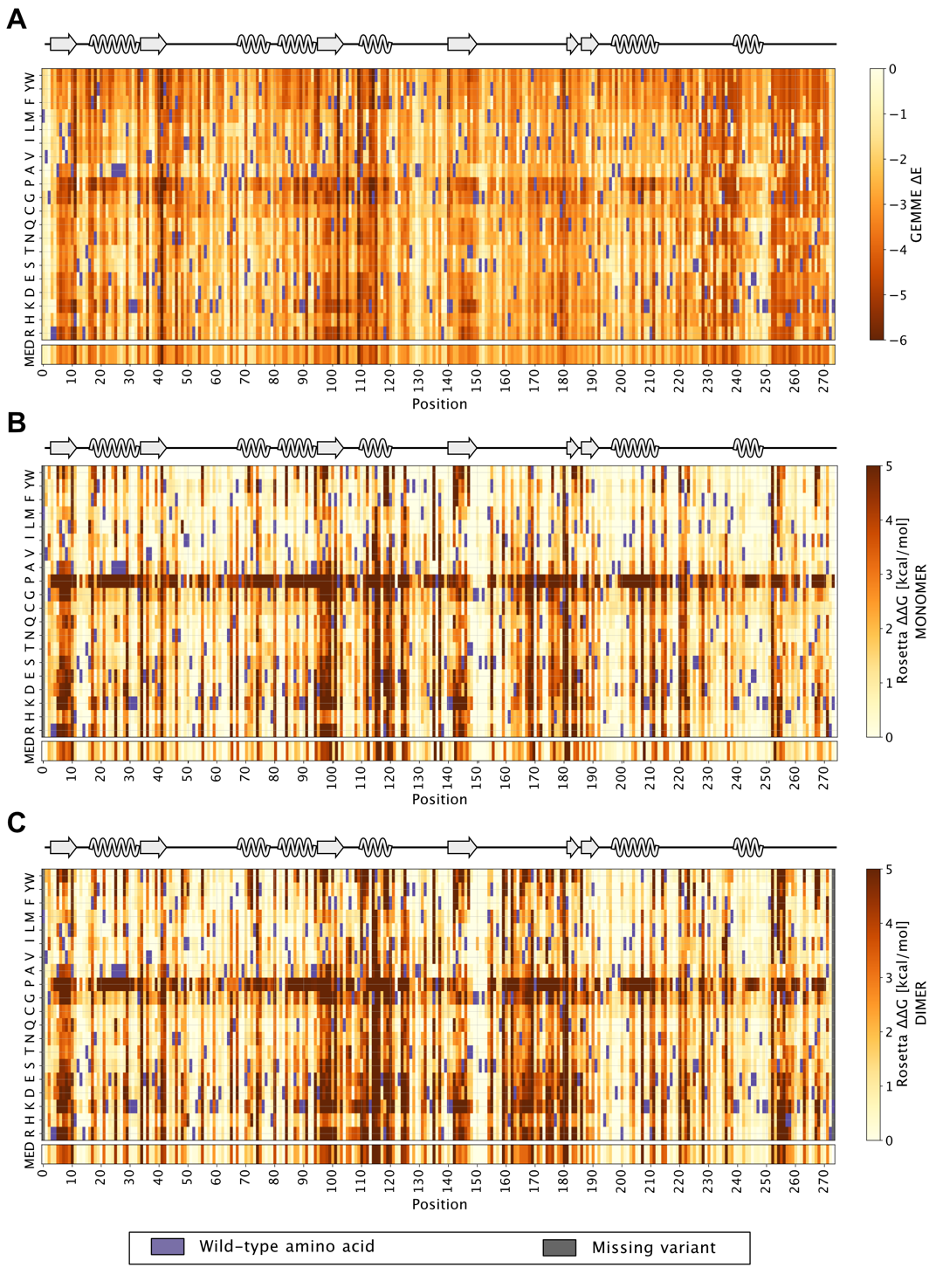

**Supplementary Figure 1. Computational prediction maps of thermodynamic stability and evolutionary conservation of NQO1.** Panel A shows the results of evolutionary conservation analysis using GEMME on NQO1 sequence. Variants evolutionary close to the WT have score close to zero, while variants with detrimental effects on the protein have high negative scores (red shadows). WT amino acid is indicated with a purple box and the median score is shown for each residue. Panel B and C shows the thermodynamic stability (ΔΔG) maps of NQO1 evaluated using Rosetta using the monomer (B) and the dimer (C) structure. Variants with a stability similar to the wild-type have scores close to zero, while variants with detrimental effects on the protein’s stability have large positive scores (coloured with shadows of red). Wild-type amino acids are indicated with purple boxed, positions (and therefore variants) which are missed in the PDB structure used are coloured in grey and the median score is shown for each position.

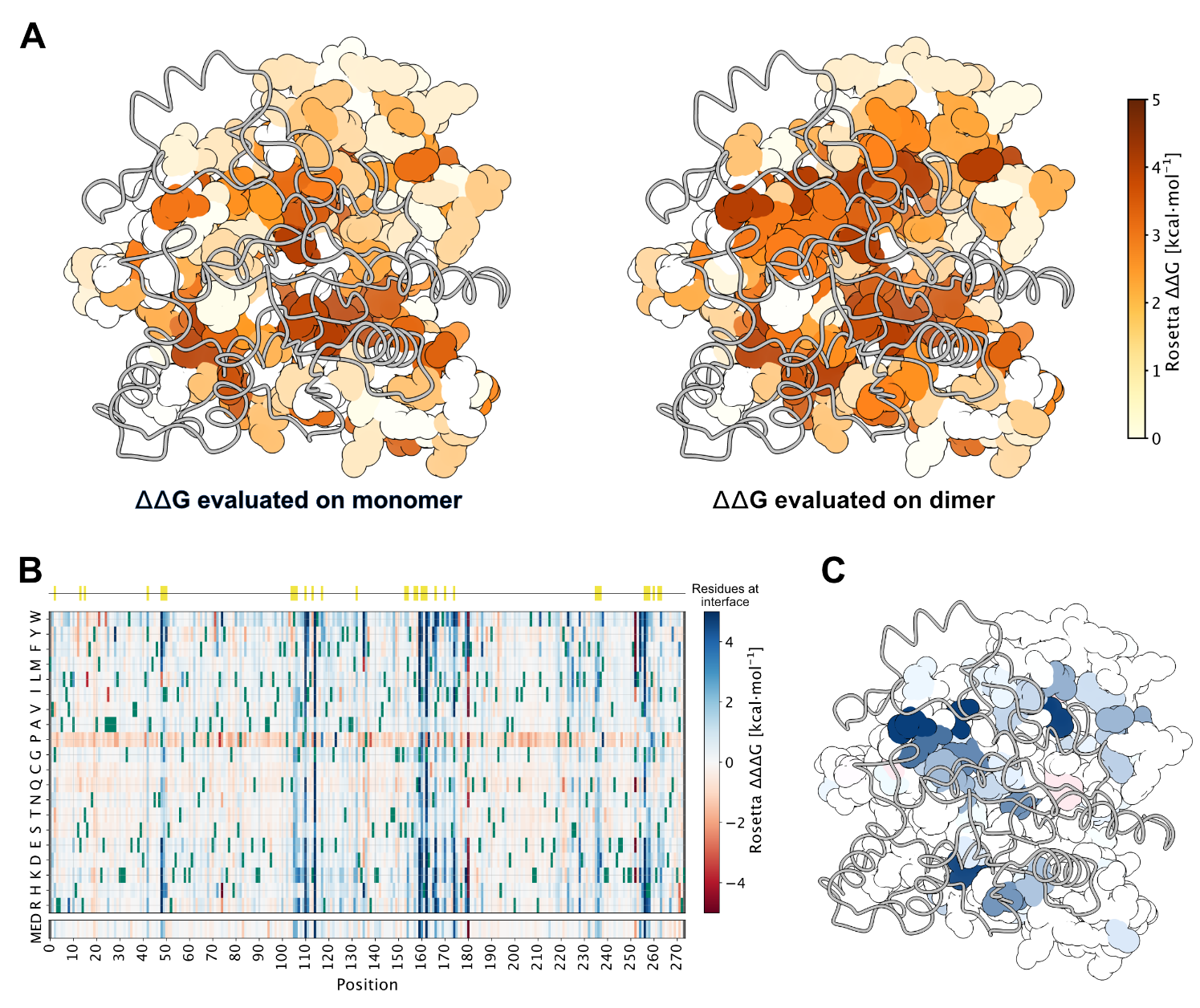

**Supplementary Figure 2.** **Thermodynamic stability differences (ΔΔG) between the monomer and dimer based on computational methods.** Panel A shows the median ΔΔG value for each residue on the crystal structure of NQO1, the figure on the left show median ΔΔGs evaluated keeping only the monomer structure of the NQO1 while the figure on the right report median ΔΔG evaluation using homozygous mutation on the dimer. Positions with neutral effect are coloured in white, while detrimental positions are coloured in shadows of red. Panel B shows the heatmap with the Δ(ΔΔG) between the two different ΔΔG evaluations on the monomer and dimer. Variants with a higher stability in dimer evaluation of ΔΔG are coloured in blue while variants with a higher stability in the monomer ΔΔG are coloured in red. WT residues are reported in green. On the top of the heatmap, interface residues are reported with a yellow marker. Panel C shows the median positional Δ(ΔΔG) difference between the two evaluations mapped to the crystal structure of NQO1.

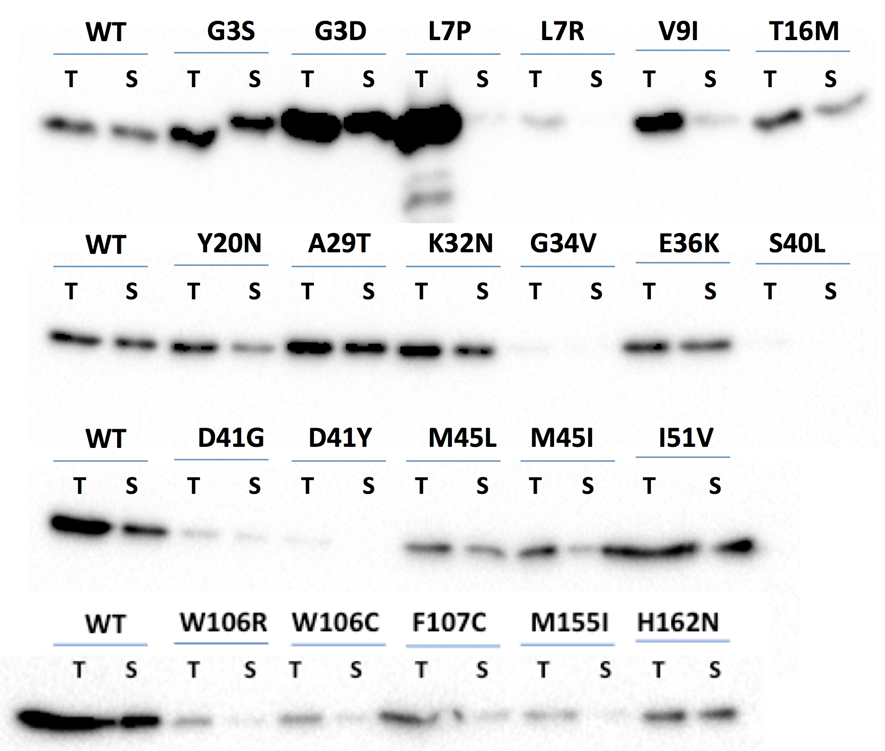

**Supplementary Figure 3. Representative Western-blot analyses of total (T) and soluble (S) expression levels for NQO1 variants in *E.coli*.** Experimental details can be found in the main text.

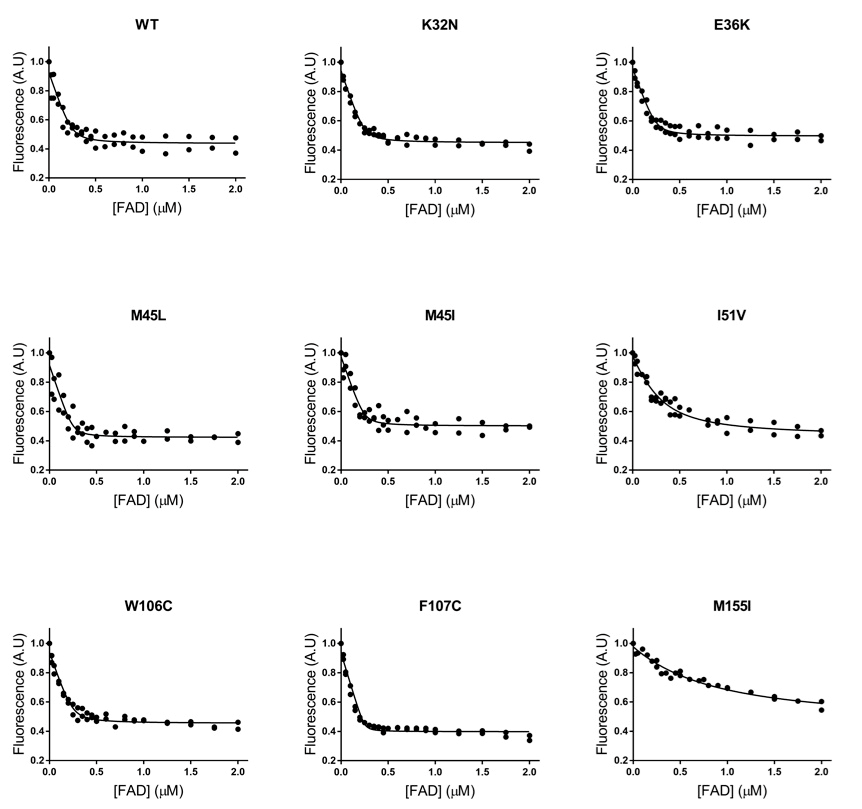

**Supplementary Figure 4. Titrations of apo-NQO1 proteins with FAD.** Experiments were replicated twice and all data were fitted to single-site binding model. In all the cases, titrations were carried out by fluorescence measurements. Experimental details can be found in the main text.

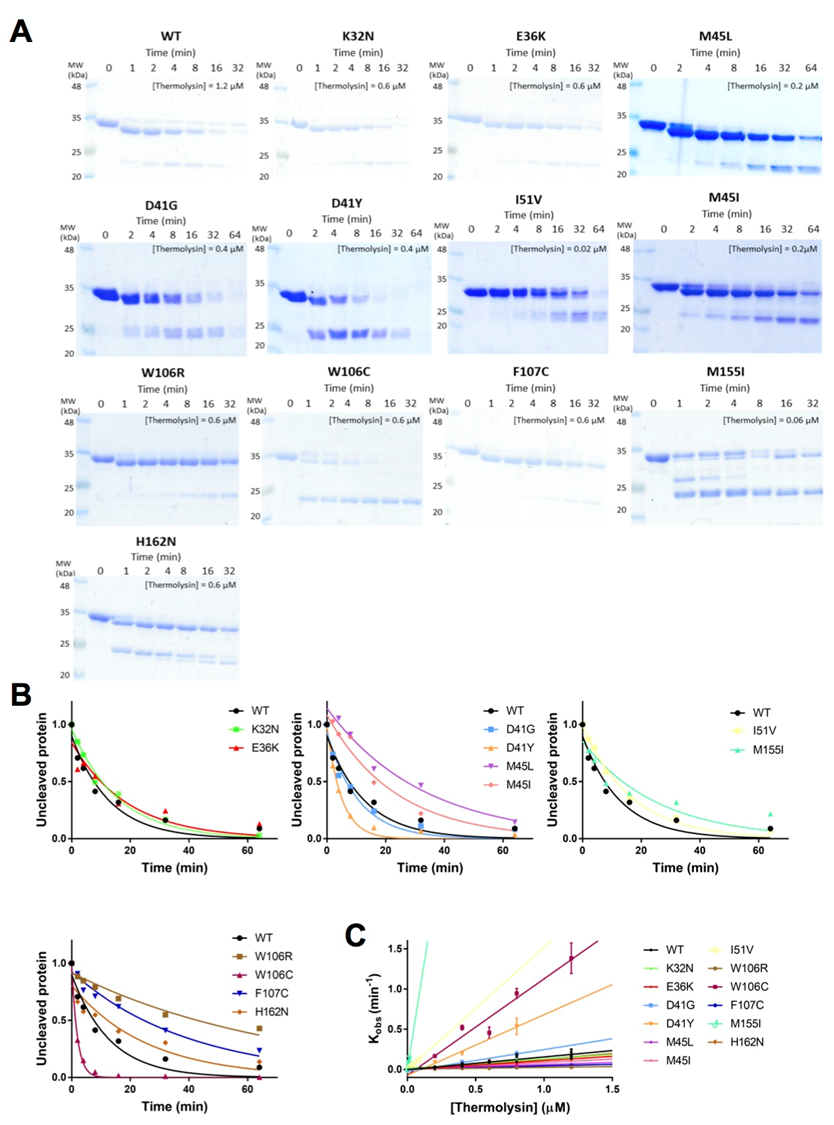

**Supplementary Figure 5. Partial proteolysis of NQO1 variants by thermolysin.** Panel A shows representative kinetic experiments with the protease and NQO1 variants. The concentration of protease used is indicated in each case. Panel B shows the corresponding densitometric analysis of native protein over time used to obtained the observed first-order rate constant using 0.4 µM thermolysin. Panel C shows the linear dependence of the observed rate constants that can be interpreted as effects on the thermodynamic stability of the TCS. Experimental details can be found in the main text.

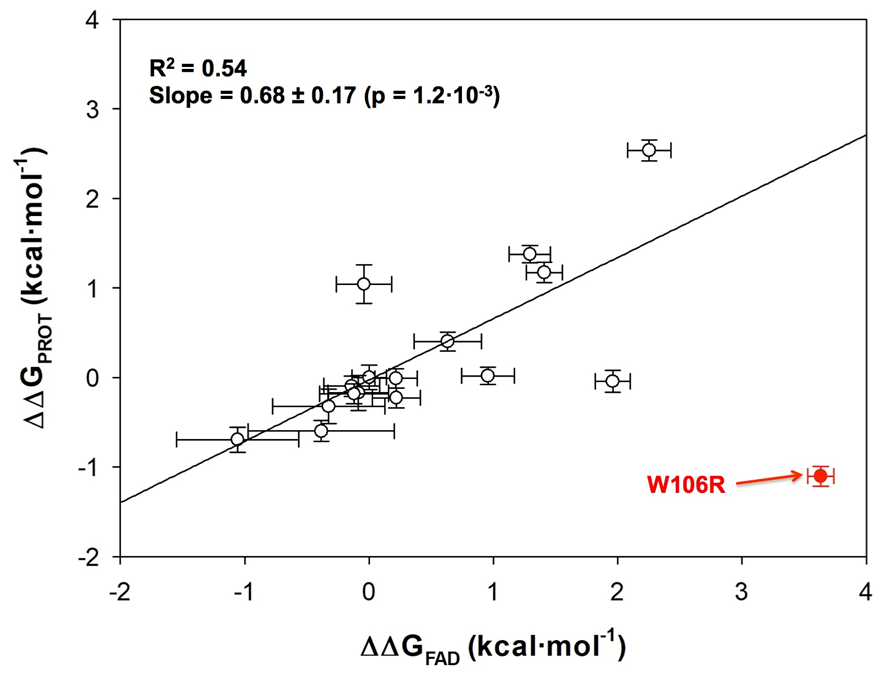

**Supplementary Figure 6. Correlation between changes in free energy for limited proteolysis (ΔΔG_PROT_)at the NTD and binding affinity for FAD (ΔΔG_FAD_).** The plot includes the best linear correlation between both parameters. W106R was excluded as a clear outlier.

**Supplementary References**

1. Pacheco-García JL, Cano-Muñoz M, Sánchez-Ramos I, Salido E, Pey AL. (2020). Naturally-Occurring Rare Mutations Cause Mild to Catastrophic Effects in the Multifunctional and Cancer-Associated NQO1 Protein. J. Pers. Med. 10, 207. doi: 10.3390/jpm10040207.
